# Small mammals and their burrows shape the distribution of *Coccidioides* in soils: a long-term ecological experiment

**DOI:** 10.1101/2024.09.21.613892

**Authors:** Jennifer R. Head, Simon K. Camponuri, Amanda K. Weaver, Lilliam Montoya, Erika Lee, Molly Radosevich, Isabel Jones, Robert Wagner, Abinash Bhattachan, Grace Campbell, Nicole Keeney, Philip A. Collender, Alexandra K. Heaney, Lisa Couper, Kate A. Colvin, Loralee Larios, William T. Bean, John Taylor, Justin V. Remais

## Abstract

**Background:** For nearly all human fungal pathogens, the environmental constraints on their distributions remain poorly understood, hindering disease management. Here, we investigated the role of zoonotic host presence, soil conditions, and their interaction on the presence of *Coccidioides immitis* – an emerging fungal pathogen and causative agent of coccidioidomycosis.

**Methods:** A long-term experimental study initiated in 2007 in the Carrizo Plain National Monument, California excluded rodents from certain areas (20 by 20-meter exclosures; n = 20) via the installation of submerged mesh fences, leaving emptied burrows intact. We collected 1,988 soil samples over four sampling periods spanning April 2021 to April 2022 using a factorial design that crossed burrows and surface soils (i.e., top 10 cm of soil) with exclosures (i.e., rodent absence) and non-exclosures (i.e., rodent presence). We measured *in situ* soil moisture, temperature, and vegetation, and analyzed soils for *C. immitis* DNA.

**Results:** The proportion of samples containing *C. immitis* was significantly higher (p <0.0001) in burrow samples taken from outside rodent exclosures (i.e., rodents present; 28.5% positive) compared to burrow samples taken from within exclosures (i.e., rodents absent; 19.7%), surface soils outside of exclosures (3.6%) and surface soils within exclosures (0.5%). In generalized linear mixed models adjusting for soil conditions, rodent presence, and sample type (burrow vs. surface soil), we estimated that the odds of detecting *C. immitis* were 19.2 (95% CI: 11.3 – 36.7, p < 0.0001) times higher in soils taken from burrows compared to surface soils, and 2.6 (95% CI: 1.0 – 5.6, p = 0.049) times higher in soils where rodents were present compared to absent. Soil moisture was the only abiotic factor associated with *C. immitis* detection (odds ratio per 1% increase in water content: 0.83, 95% CI: 0.70 – 0.89, p = 0.0002). In mediation analyses, we estimated that 73.7% (95% CI: 68.9, 78.5) of the total association between rodents and *Coccidioides* detection effect was attributable to rodent creation of burrows.

**Conclusions:** Rodent creation and occupation of burrows substantially increases *C. immitis* detection. Synergy between reservoir host presence and environmental conditions may exist for other human fungal pathogens.

## Introduction

Fungal pathogens pose a large and increasing threat to global public health, with over 1.5 million deaths annually, rising antifungal resistance and no antifungal vaccines, and, in many cases, expanding geographic distributions (Nnadi and Carter 2021, Seagle, Williams and Chiller 2021, Rokas 2022, Xu 2022). Despite this, fungi remain underrecognized as pathogens of human concern, and knowledge gaps persist in understanding the ecological drivers of their maintenance and spread (Rodrigues and Nosanchuk 2021). Human-pathogenic fungi can be classified according to transmission route (human-to-human vs. acquired from the environment) and host type (primary pathogens that establish infection in healthy hosts vs. opportunistic which require an immunocompromised host). The prevention of environmentally acquired, primary fungal infections requires comprehensive understanding of their ecology, including the factors affecting where they grow in the environment. While most fungal pathogens exhibit broad host ranges and have high dispersal capacity, primary environmental fungal pathogens have more narrow geographic ranges (Fisher, Henk et al. 2012). Adaptation to particular environmental conditions or hosts may play a key role in constraining the geographic distribution of environmental fungal pathogens (Fisher, Henk et al. 2012). As animals have coevolved with fungi, the pathogenicity of human-pathogenic fungi is thought to have evolved for infecting specific animals before infecting humans (Köhler, Casadevall and Perfect 2014). The environmental distributions of several primary environmental fungi are associated with presence of certain zoonotic hosts: *Talaromyces marneffei* is associated with bamboo rats (Mahajan 2021), *Histoplasma capsulatum* growth is promoted by guano from bats as well as chickens and other birds (Benedict and Mody 2016), and armadillos are considered a natural reservoir for *Paracoccidioides* spp.,(Martinez 2017). However, the relative contribution of animal hosts versus host habitats (and the associated soil conditions within these habitats) on their distribution remain unknown (Větrovský, Kohout et al. 2019), hindering disease control.

The role of zoonotic hosts in the life cycle and distribution of *Coccidioides –* another environmentally-acquired, primary fungal pathogen of public health importance – has also remained unresolved. *Coccidioides* is a soil fungus endemic to desert-like regions of North and South America and a causative agent of coccidioidomycosis, an emerging fungal disease in southwestern U.S. (Barker, Litvintseva et al. 2019, Taylor and Barker 2019, Kirkland, Stevens et al. 2022). *Coccidioides* is known to infect a variety of small mammals, and particularly burrowing rodents (Emmons and Ashburn 1942, Emmons 1943, Barker 2018, Salazar-Hamm, Montoya et al. 2022) including kangaroo rats, pocket mice (Emmons and Ashburn 1942, Emmons 1943) deer mice, and woodrats (Catalan-Dibene, Johnson et al. 2014). In the environment, *Coccidioides* is commonly found in and around rodent burrows within the endemic region (Barker, Tabor et al. 2012, Kollath, Teixeira et al. 2020), as well as in soils near carcasses of infected mammals (Maddy and Crecelius 1965). *Coccidioides* growth in the soil is facilitated by moisture and organic matter (Maddy 1957), with comparative genome analyses demonstrating that *Coccidioides* has lost genes associated with the breakdown of plant cell wells, yet harbors an abundance of genes associated with digestion of animal protein (Sharpton, Stajich et al. 2009). These lines of evidence have led to the hypothesis that burrowing mammals are reservoir hosts of *Coccidioides* spp., harboring fungal spores without experiencing disease, and, upon death, providing a substrate for *Coccidioides* growth in soils (Swatek and Plunkett 1957, Del Rocío Reyes-Montes, Pérez-Huitrón et al. 2016, Taylor and Barker 2019), This hypothesis is termed the ‘endozoan small-mammal reservoir hypothesis’.

However, this hypothesis is the subject of active debate, with an alternative theory proposing that burrows themselves may provide beneficial conditions that support *Coccidioides* populations, independent of rodent occupation of burrows (Lacy and Swatek 1974, Del Rocío Reyes-Montes, Pérez-Huitrón et al. 2016) Rodent burrows may provide favorable microhabitat conditions including relatively higher soil moisture, lower thermal fluctuations, decreased soil bulk density, and altered vegetation coverage (Studier and Baca 1968, Mun and Whitford 1990, Whitford and Kay 1999, Gurney, Prugh and Brashares 2015, Mead, Hamm et al. 2020). Prior attempts to define the ecological niche of *Coccidioides* have often found associations with hot, arid, alkaline soils (Baptista-Rosas, Hinojosa and Riquelme 2007, Fisher, Bultman et al. 2007, Weaver and Kolivras 2018, Ocampo-Chavira, Eaton-Gonzalez and Riquelme 2020). The extent to which these microhabitat conditions alone promote *Coccidioides* maintenance and growth, versus the host presence itself, has remained unclear.

Long term ecological experiments are invaluable in estimating how species distributions respond to changes in experimentally manipulated abiotic and biotic conditions. Here, we leveraged a long-term experimental rodent-exclosure study to resolve the role of zoonotic host reservoirs, environmental conditions, and their interaction in the lifecycle and distribution of *Coccidioides immitis*, a species of *Coccidioides* endemic to California. The results provide key insights into the endozoan small-mammal reservoir hypothesis as well as evidence of interaction between environmental conditions and animal reservoir hosts that may aid in understanding the environmental distribution of other primary environmental fungal pathogens.

## Methods

### Study site

The Carrizo Plain National Monument in San Luis Obispo County, California, is the largest intact desert-grassland within California, containing annual grassland and alkali desert scrub habitats (Endicott, Dillard et al. 2017). It is home to several species of kangaroo rat that have previously been identified as hosts for *Coccidioides* (Emmons and Ashburn 1942, Emmons 1943). The largest, the giant kangaroo rat (*Dipodomys ingens*), are considered ecosystem engineers because they construct burrows known as precincts, which are used by other animals and form distinctive mounds on landscapes (Figure 1A)(Corten 2008). In normal conditions, the giant kangaroo rat excludes or greatly outnumbers other nocturnal rodent species, although during droughts, smaller-bodied rodents increase in relative abundance as giant kangaroo rat populations decline (Prugh, Deguines et al. 2018). The California Carrizo Plains and its surrounding areas are endemic for *Coccidioides* (Lauer, Lopez et al. 2023) and outbreaks have occurred in and near the plain (Wilken, Sondermeyer et al. 2015).

**Figure 1.**
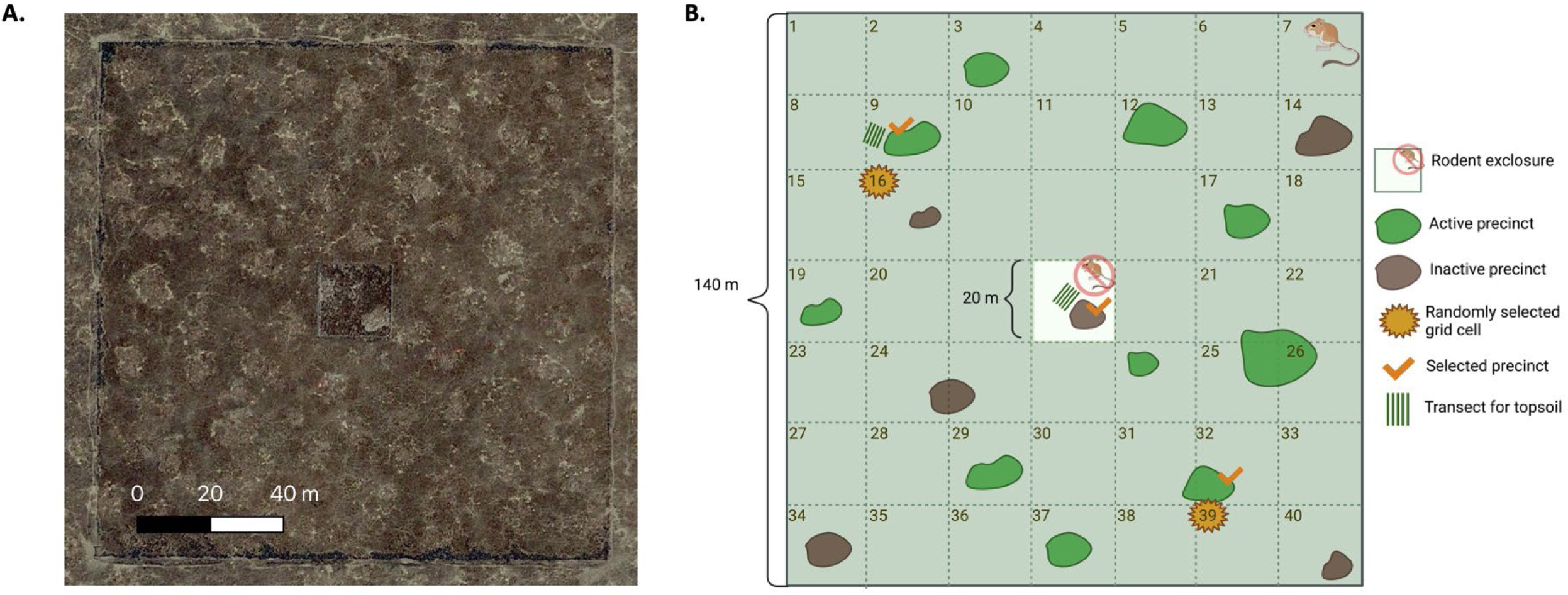
Satellite image (A.) and illustrative schematic (B.) of one experimental plot within the Carrizo Plain Ecosystem Project (CPEP) experimental study. Each plot is 140m x 140m with a 20 x 20 m rodent exclosure in the middle. A) Satellite image of an experimental plot, obtained from Google Satellite Imagery. Light colored patches are precincts, and the square in the center is the exclosure. The plot perimeter is fenced to exclude cattle. B.) Each plot was divided into grids, and two grid cells outside of the exclosure were selected for sampling using a random number generator (cells 16 and 39 in this schematic). One precinct in the exclosure was sampled, as was the active precinct nearest to each selected non-exclosure grid cell. Five surface soil samples were collected from two transects radiating 2, 3, 4, 5, and 6 meters away from a single precinct in the exclosure and non-exclosure.

A long-term, experimental study—the Carrizo Plain Ecosystem Project (CPEP)—was launched in 2007 to study the relationships between giant kangaroo rats, cattle, vegetation, and other species using replicated exclosures (Endicott, Dillard et al. 2017). Twenty experimental plots are split between two pastures; while cattle are excluded from all plots, the northern pasture has permitted historical cattle grazing. Each plot is 140 x 140 square meters and contains a 20 x 20 meter rodent exclosure in the middle (Figure 1). The exclosure is a submerged fence of hardwire cloth (0.25 inch mesh) that extends two feet above ground and two feet below ground, to prevent burrowing rodents from entering (Prugh and Brasheres 2007). At the time of exclosure creation, giant kangaroo rats found inside the fence were trapped and relocated outside the exclosure. The exclosure encompasses precincts occupied prior to 2007, from which rodent access has been restricted for 15 years. Each spring, giant kangaroo rats that have breached the exclosure are trapped and relocated. Breaching of the exclosure by smaller rodents – typically by the San Joaquin antelope squirrel – occurs occasionally; these mammals have minimal engineering effects (e.g., altering vegetation, building burrows) and are unlikely to establish in the exclosure.

### Environmental sampling

Across four sampling seasons (April 2021, July 2021, October 2021, and April 2022), we collected soil samples within 20 CPEP experimental plots, split evenly by pasture (Figure 2). Within each plot, we collected samples across a factorial design of burrows and surface soils crossed with exclosures and non-exclosures, for a total of four sampling combinations. During each season, a target sample size of 25 soil samples were collected per plot, consisting of five burrow and five surface soil samples within the exclosure, and 10 burrow and five surface soil samples outside of the exclosure (Figure 1B). The larger number of samples collected from burrows outside the exclosure was chosen to maximize statistical power, on the basis of sample size calculations. Thus, our total target sample size was 500 in each of the four sampling seasons.

**Figure 2.**
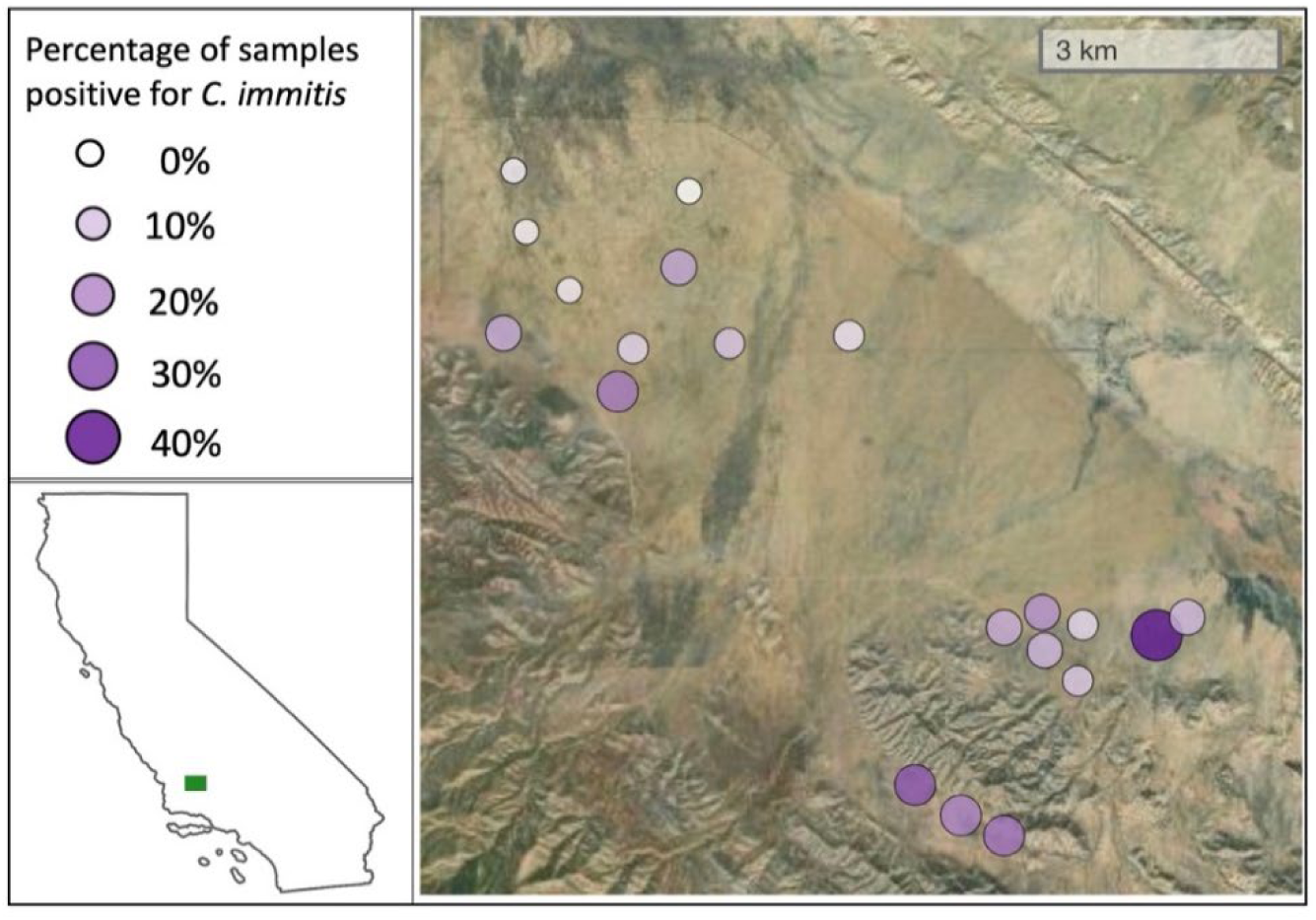
Percentage (%) of samples per plot that tested positive for Coccidioides in soils. Each symbol represents all samples drawn from one experimental plot in the Carrizo Plain Ecosystem Project, pooled across seasons (n = ∼25 samples per season, totaling approximately 100 per plot across four seasons). Inset map indicates location of the Carrizo Plain, in green. Maps stratified by seasons and sample type are included as Supplemental Figures 4-5.

Within each exclosure, we sampled soils from five unique precinct burrow entrances. Burrow entrances (burrows hereafter) were sampled by excavating approximately 10 mL of soil from burrow walls approximately 20-30 cm from the burrow entrance using a coffee scoop with a ten-inch handle. In some seasons we could not identify five unique precinct entrances within certain plot exclosures, and thus collected fewer than 500 samples that season. In total, we collected and analyzed 1,988 soil samples for the presence of *Coccidioides* (Table 1). Surface soils were sampled along a linear transect that radiated outward from the fifth sampled burrow opening, with the first sample collected two meters from a burrow opening, and the remaining four samples spaced one meter apart. Surface soil was collected by digging a hole ten centimeters deep with a trowel and sampling from the walls and bottom of the hole. Ten centimeters was chosen as the sampling depth in order to capture the area of soils where *Coccidioides* is thought to be most likely to occur (Fisher, Bultman et al. 2007), and to roughly approximate the vertical depths that soils collected from within burrows were sampled from. Within the exclosure, burrows that seemed active indicating a potential breach were not sampled, nor were burrows that were within two meters of the exclosure fence. Signs of rodent activity included tail or feet markings around burrow entrance, clipped vegetation or seed casings near burrow entrance, or fecal pellets (Figure S1).

**Table 1.** Prevalence of *Coccidioides* detection, stratified by sampling season and sample type.

| Positive samples / Total samples (% positive) |  |  |  |  |  |
| --- | --- | --- | --- | --- | --- |
|  | Burrows |  | Surface soils |  | Total |
|  | Rodents excluded | Rodents present | Rodents excluded | Rodents present |  |
| **Spring 2021** | 12/99 (12.1%) | 47/200 (23.5%) | 0/100 (0%) | 4/99 (4.0%) | 63/498 (12.7%) |
| **Summer 2021** | 21/97 (21.6%) | 76/200 (38%) | 0/99 (0%) | 2/100 (2.0%) | 99/496 (20.0%) |
| **Fall 2021** | 27/100 (27.0%) | 65/200 (32.5%) | 0/100 (0%) | 3/95 (3.2%) | 95/495 (19.2%) |
| **Spring 2022** | 18/100 (18.0%) | 40/199 (20.1%) | 2/100 (2.0%) | 5/100 (5.0%) | 65/499 (13.0%) |
| **Total** | 78/396 (19.7%) | 228/799 (28.5%) | 2/399 (0.5%) | 14/394 (3.6%) | 322/1988 (16.2%) |

Outside of exclosures, we identified the precincts and associated surface soils to sample using a systematic, random sampling procedure established during our first sampling period (April 2021). First, we divided the plot into a 7 x 7 grid, excluding grid cells contiguous with the exclosure (Figure 1B), and selected two non-contiguous cells using a random number generator. From each selected grid cell, we located the closest precinct that had signs of rodent activity. For the precinct furthest from the exclosure, we collected surface soil samples as described previously. Sampled precincts and the surface soil transect direction were marked in April 2021 with flags, and GPS coordinates were logged, allowing sampling from the same precincts in subsequent seasons. In a few cases, flags could not be identified during return trips, and we sampled from the precinct that most closely matched the recorded geolocation.

All sampling tools were cleaned with 70% ethanol between individual sample collections. For each sample, we recorded information on soil moisture (measured as percent volumetric water content; Campbell Scientific® HS2 probe), soil temperature, and vegetation (type: barren, grass or shrub; level: none = 0, low = 1, medium = 2, high = 3). Data were collected using REDcap® mobile survey platform (UCSF). For each of the four sample types in a single plot, we collected a continuous measurement of temperature over a 24- to 48-hour period using temperature loggers placed within the burrows or buried ten centimeters beneath the surface (HOBO® MX2202).

### Laboratory analysis of soils

Soil samples were stored in centrifuge tubes at room temperature until analysis, following standard practice (Johnson, Carlson et al. 2014, Alvarado, Teixeira et al. 2018, Wagner, Montoya et al. 2022, Wagner, Montoya et al. 2023). Prior work demonstrates that *Coccidioides* survival and meta-barcoding of soil fungi is similar at room temperature, 4°C, and -15°C (Friedman, Smith et al. 1956, Delavaux, Bever et al. 2020). We extracted DNA using the Qiagen PowerSoil kit, following established protocol (Greene, Koenig et al. 2000, Barker, Tabor et al. 2012, Kollath, Teixeira et al. 2020, Wagner, Montoya et al. 2022), and used the CocciEnv qPCR assay (Bowers, Parise et al. 2018) to determine presence of *C. immitis* in the sample. Details of the protocol are included in Supplemental Methods. We ran four replicates of each DNA aliquot and classified a sample as positive for *Coccidioides* if at least three of four wells had a cycle threshold value below 40. We conducted sensitivity analyses that varied this criteria (Supplemental Methods & Results).

### Statistical analysis

#### Generalized liner mixed models

We fit generalized linear mixed models (GLMMs) to estimate associations between rodents, burrows, soil conditions, and *C. immitis* detection. We regressed the binary outcome (*Coccidioides* presence) against binary indicators for rodent presence (vs. exclusion), burrow (vs. surface soil), pasture (northern or southern), and indicators for sampling trip. To account for the clustered sampling design and repeated measures, we included random effects representing unique sampling clusters in the model, nested within random effects on plot. We denoted three clusters per plot: the paired precinct and surface soil transect exclosure samples (n=10), the paired precinct and surface soil transect non-exclosure samples (n=10), and the unpaired precinct non-exclosure samples (n=5). In total, we had 60 random effects (three per plot nested within 20 plots). We built separate models that did and did not control for soil moisture (continuous), vegetation level (ordinal, regardless of type), and soil temperature (continuous) as fixed effects.

#### Causal mediation analysis

We sought to disentangle the impacts of rodent presence, burrows, and soil conditions on the probability of *Coccidioides* detection. We conducted a causal mediation analysis with g-computation (Wang and Arah 2015), which allows us to compute the percent difference in *Coccidioides* detection attributable to each of these factors (Figure 3). In this framework, rodents are the binary exposure of interest (*X*; 1 = present, 0 = absent), burrows the mediator (*M*; 1 = burrow, 0 = surface soil), and *Coccidioides* presence in soil the outcome (*Y*) (Supplemental Methods). Where used, soil conditions (moisture, temperature, vegetation) are represented among the set of covariates (***Z***), along with sampling pasture and season. In sensitivity analyses, we repeated analyses, dropping plots where concern of giant kangaroo rat breach had been noted across more than one sampling season (Supplemental Methods & Results). We used a similar mediation modeling framework to examine the degree to which the association between burrows and *Coccidioides* presence could be mediated by soil conditions (Supplemental Methods).

**Figure 3.**
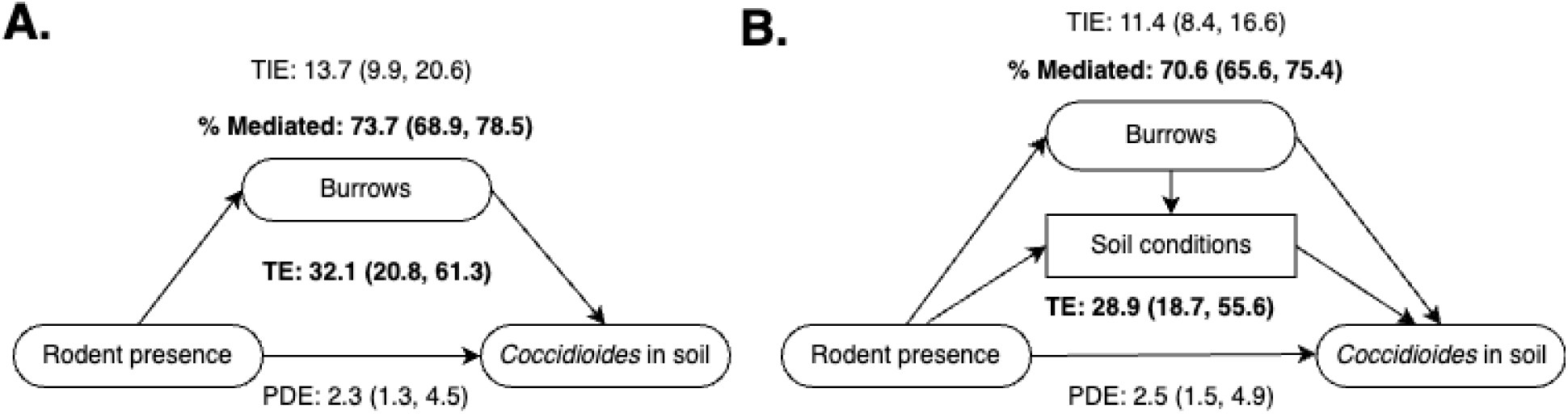
Directed acyclic graphs (DAGs) representing the hypothesized relationships between rodents, burrows, and *Coccidioides* presence in soils. Values reflect the odds ratios from causal mediation analysis not controlling for soil moisture, temperature, and vegetation (A), and controlling for these variables (B). TE = total effect; PDE = pure direct effect; TIE = total indirect effect.

## Results

*Coccidioides immitis* was detected in 322 (16.5%) of 1,988 collected samples (Table 1). The proportion of samples where *Coccidioides* was detected varied across season (Table 1; Figure S2) and was highest among soils collected from burrows outside of exclosures (228/799; 28.5%), followed by burrows within exclosures (78/396; 19.7%), surface soils outside of exclosures (13/394; 3.6%), and surface soils within exclosures (2/396; 0.5%; Table 1). Plots in the southern pasture (no historical cattle grazing) exhibited significantly higher rates of *Coccidioides* detection (p = 0.0008) as compared to plots in the northern pasture (historical cattle grazing; Figure 2; Figures S3-4). Clustering of positive samples within precincts or transects was observed for both surface soils and burrows (Figure S5; Supplemental Results), but there was no association between positivity among the transects and positivity in the precinct.

For all samples, the average soil moisture was 3.3% (standard deviation [sd]: 2.7%, range: 0% - 20.9%), the average soil temperature was 23.9°C (sd: 7.0 °C, range: -13 – 43 °C), and the most common vegetation level and type was low grass (mean level = 1; range: none [0] to high [3]). These values varied across season and sample type (Table S1; Figure S6).

### Contribution of rodents and burrows to Coccidioides presence in soil: results from GLMMs

GLMM results indicated strong, independent associations between both burrows and rodent presence with odds of detecting *Coccidioides* in the soil (Table 2). Controlling for rodent presence, season, and soil moisture, the odds of *Coccidioides* detection in soils were 19.2 times higher (95% CI: 11.3 – 36.7; p <0.0001) among soils collected from burrows than in soils collected from the top 10 cm of the surface. The magnitude of the effect of rodent presence on *Coccidioides* presence was smaller but still significant: controlling for sample type (burrow vs. surface soil), season, and soil moisture, the odds of *Coccidioides* detection in areas where rodents were present were 2.6 times higher (95% CI: 1.0 – 5.6; p = 0.049) than in soils collected in areas where rodents were excluded (Table 2). Lower soil moisture was associated with higher odds of *Coccidioides* detection, controlling for season, rodent presence, and soil type. A one percentage point increase in volumetric water content was associated with a 17% reduction in the odds of *Coccidioides* detection (OR: 0.83, 95% CI: 0.70 – 0.89; p = 0.0002). *Coccidioides* detection was significantly more frequent among samples collected during the summer (p = 0.0006) and fall (p = 0.0008) 2021 field expeditions than among those collected during the springs of 2021 and 2022. There was no effect of soil vegetation level at sample location or soil temperature on *Coccidioides* detection, nor were these factors important confounders. These variables were dropped from further analyses (Table 2).

**Table 2.** Odds ratios of *Coccidioides* detection obtained from generalized linear mixed models, *E*(*Y*|*X* = *x*, *M* = *m*, ***Z*** = ***z***). Fixed effects are as shown in the table; random effects are nested plot and paired cluster; *Y* = *Coccidioides* detection (binary); *X* = rodents; *M* = sample type (burrow vs. surface soil); ***Z*** = (season, pasture, soil moisture).

|  | Model without soil covariates |  | Model with soil covariates |  |
| --- | --- | --- | --- | --- |
|  | OR (95% CI) | p-value | OR (95% CI) | p-value |
| **Rodents present** (*ref: exclosure*) | 2.4 (0.85, 5.3) | 0.098 | 2.6 (1.0, 5.6) | 0.049 |
| **Burrows** (*ref: surface soil*) | 23.0 (14.0, 44.7) | <0.0001 | 19.2 (11.3, 36.7) | <0.0001 |
| **Soil moisture** (water content %) | -- | -- | 0.83 (0.70, 0.89) | 0.0002 |
| **Season** (reference = spring 2021) |  |  |  |  |
| Summer 2021 | 2.2 (1.5, 3.5) | 0.0002 | 1.6 (0.92, 2.4) | 0.10 |
| Fall 2021 | 2.0 (1.4, 3.4) | 0.0004 | 1.4 (0.85, 2.3) | 0.19 |
| Spring 2022 | 1.1 (0.69, 1.7) | 0.70 | 1.2 (0.80, 2.1) | 0.29 |
| Log likelihood | -631 |  | -626 |  |
| AIC | 1,278 |  | 1,270 |  |
| BIC | 1,323 |  | 1,320 |  |
| Number of observations | 1,788 |  | 1,788 |  |

### Contribution of rodents and burrows to Coccidioides presence in soil: results from causal mediation analysis

In mediation analyses that did not control for soil conditions, the odds ratio estimating the total effect (TE) of rodent presence on *Coccidioides* in soil was 32.1 (95% CI: 20.8, 61.3) indicating the odds of detecting *Coccidioides* in an area with current, active rodent presence is over 30 times higher than in the counterfactual scenario where rodents have never been present in the sampled location (Figure 3A, Table 3). This estimate captures the effect of rodents on *Coccidioides* via both indirect effects (e.g., building burrows that support *Coccidioides* presence, modifying local soil conditions) and direct effects (e.g., increasing the bank of arthroconidia (the infectious life stage) in the soil by providing protein and water for hyphal growth and sporulation, and releasing spores from recently deceased rodent carcasses).

**Table 3.** Odds ratios representing various effects from causal mediation analysis examining relationship between rodents, burrows, soil conditions, and *C. immitis* in soils.

| Research question, adapted from [(Wang and Arah 2015)] | Effect and counterfactual definition* | Set of Y simulated using: | Estimate, unadjusted for soil conditions (Figure 3A)<br>OR (95% CI) | Estimate, adjusting for soil conditions (Figure 3B)<br>OR (95% CI) |
| --- | --- | --- | --- | --- |
| Overall, to what extent (relative) does contemporary rodent presence (*X*) increase probability of *Coccidioides* detection (*Y*) compared to the counterfactual scenario where rodents have never been present? | Total effect (TE)<br>$E[Y_{x1m_{x1}} - Y_{x0m_{x0}}]$ | *x* simulated according to observed distribution. *m<sub>x</sub>* is simulated using *x* values that follow the observed distribution. | 32.1 (20.8, 61.3) | 28.9 (af) |
| By what factor does rodent presence (*X*) versus absence amplify *Coccidioides* presence (*Y*), through pathways other than via burrows (*M*)? | Pure direct effect (PDE)<br>$E[Y_{x1m_{x0}} - Y_{x0m_{x0}}]$ | *x* simulated according to observed distribution. *m<sub>x</sub>* is simulated using *x* set to 0 (*m<sub>x0</sub>*). | 2.3 (1.3, 4.5) | 2.5 (1.5, 4.9) |
| When rodents are present (*X* = 1), by what factor does burrow creation (*M*) by rodents amplify *Coccidioides* detection (*Y*)? | Total indirect effect (TIE)<br>$E[Y_{x1m_{x1}} - Y_{x1m_{x0}}]$ | *x* is set to 1 (*x*<sub>1</sub>). *m<sub>x</sub>* is simulated using *x* values that follow the observed distribution. | 13.7 (9.9, 20.6) | 11.4 (8.4, 16.6) |
| What proportion of the effect of rodents (*X*) on *Coccidioides* (*Y*) is mediated by burrows (*M*)? | Percent of effect mediated via burrow (%) |  | 73.7 (68.9, 78.5) | 70.6 (65.6, 75.4) |

\*Y outcome (*Coccidioides* detection), *X* exposure (rodents present), *M* mediator (burrow). Subscripts *x*<sub>1</sub> and *x*<sub>0</sub> represent the index and counterfactual values of the exposure (rodents currently present vs. rodents never present) and *m<sub>x1</sub>* and *m<sub>x0</sub>* represent the expected value of the mediator (burrows) when the exposure is set to the index or the counterfactual. As described in methods, *m<sub>x1</sub>* = 0.5 and *m<sub>x1</sub>* = 0.

The direct effect (see Supplemental Methods) of rodents on *Coccidioides* detection was 2.3 (95% CI: 1.3, 4.5), representing the degree to which rodents increase the odds of detecting *Coccidioides* through pathways other than burrow creation. In other words, the odds of detecting *Coccidioides* in a counterfactual world where rodents are present but do not burrow is 2.3 times higher than the odds of detecting it in a counterfactual world where rodents are absent. The indirect effect of rodents on *Coccidioides* that operates via burrows was 13.7 (95% CI: 9.9, 20.6). This means that, in a world (such as the one that exists!) where rodents are present and able to create burrows, the odds of *Coccidioides* detection was 13.7 times higher than in a counterfactual world where rodents are present but do not burrow. The percent of the total effect that is mediated is equal to the log of the TIE divided by the log of the TE. We estimated that 73.7% (95% CI: 68.9, 78.5) of the effect of rodents on *Coccidioides* presence was attributable to burrow creation (Figure 3A, Table 3).

Controlling for soil moisture resulted in similar results as mediation analyses that did not include soil moisture, with a slight attenuation of the estimated total (TE: 28.9, 95% CI: 18.7, 55.6) and indirect (TIE: 11.4, 95% CI: 8.4, 16.6) effects (Figure 3B; Table 3). This small attenuation implies that changes to soil conditions induced by rodent and burrow presence accounted for a small amount of the effect of rodents on *Coccidioides* presence. After accounting for the effect of soil conditions, the percent of the total effect of rodent presence on *Coccidioides* that was mediated by burrows was slightly lower (71.7%; 95% CI: 67.6, 76.5%).

### Role of soil conditions in Coccidioides presence, and role of soil conditions as a mediator

Soil moisture was significantly lower among burrows compared to surface soils and was unassociated with rodent presence (Figure S6; Table S1). Controlling for season, volumetric water content, was, on average, 1.4 percentage points lower in burrows compared to surface soils (95% CI: 1.2 – 1.6, p = 0.0008). In mediation analyses to understand how much of the effect of burrows on *Coccidioides* could be mediated by lower soil moisture in burrows, we found that 8.1% (95% CI: 4.6, 11.3) of the total effect of burrows on *Coccidioides* was mediated by altered soil moisture within burrows (Figure S7; Table S2).

Temperature and vegetation differed across sample type as well but were not associated with *Coccidioides* presence in GLMMs. Continuous, overnight measurements of soil temperature revealed that surface soils experienced greater temperature fluctuations than burrows (Figure S8). At night, when temperatures were cooler, burrows were warmer than surface soils; during the day, when temperatures were hot, burrows were cooler than surface soils (Figure S8). Rodent presence was associated with lower vegetation by roughly three-fourths of an ordinal level (e.g., from medium to low; model coefficient: -0.76, 95% CI: -0.56 – -0.97), while burrows were associated with slightly higher vegetation (coefficient: 0.18, 95% CI: 0.12 – 0.24).

## Discussion

The importance of zoonotic hosts and microhabitat conditions on the distribution and spread of *Coccidioides* is recognized as one of the key outstanding questions in the field (Gorris, Caballero Van Dyke et al. 2021). We found evidence that both rodent presence and their burrows (occupied or not) are independently associated with increased probability of detection of *C. immitis* in soils. Using a carefully controlled experimental design, we found a reduction in probability of presence of *C. immitis* in burrows from 28.5% to 19.7% following 15 years of rodent exclusion. Using a causal mediation approach, we estimated that the odds of detecting *Coccidioides* in an area where rodents are currently present was nearly 32 times higher than in the counterfactual scenario where rodents have never been present. While we estimated that most of this effect was mediated via burrow creation, 26-28% of the association between rodent presence and *Coccidioides* detection was direct, suggesting a mechanistic relationship between rodent presence and *Coccidioides* distribution that is attributable to factors other than burrow creation.

Several hypotheses may explain our finding of a direct association between rodent presence and *Coccidioides* detection. One hypothesis – the ‘endozoan small-mammal reservoir hypothesis’ – asserts that *Coccidioides* live in granulomas formed within rodent hosts. Upon host death, the fungi are released from the granulomas, and use the rodent carcass as a substrate for growth, and may then go on to infect other hosts, perpetuating the cycle (Taylor and Barker 2019). Another explanation is that burrowing mammal presence concentrates nutrients like keratin (from rodent fur, skin, and nails), nitrogen, or calcium in the soil, supporting fungal growth (Gurney, Prugh and Brashares 2015). While many fungi are capable of degrading keratin, genes for doing so are more abundant in *Coccidioides* spp. and other members of the Onygenaceae family compared to other fungi families (Sharpton, Stajich et al. 2009). In our GLMMs, *Coccidioides* presence was unassociated with vegetation levels, supporting the idea that the fungus degrades nutrients other than plant materials. However, as this study did not involve investigating *Coccidioides* infection rates in small mammal species or measuring soil nutrient profiles, we were unable to disentangle these underlying mechanisms – doing so remains an important future research direction.

The strong indirect effects of burrows on *Coccidioides* presence that we detected suggests that the burrow environment may serve as a suitable microhabitat for the fungus, regardless of rodent presence. We detected *Coccidioides* in nearly 20% of burrow samples taken from where rodents had been excluded for at least 15 years. While *Coccidioides* was notably higher in rodent-occupied burrows (28.5%), the presence of pathogen populations in burrow soils long since abandoned by active rodent populations nonetheless suggests that rodent removal alone may not be a sufficient control strategy. However, we are unable to determine if presence of *Coccidioides* within burrows is due to active growth within the burrows versus entrapment of spores blown there during wind erosion or long-term persistence of inactive spores. *Coccidioides* spores can survive at least ten years in the soil (Maddy and Crecelius 1965, Chow, Kangiser et al. 2021), and it is feasible that effects of historical rodent presence (e.g., increased keratin in soil, old carcasses) are still contributing to *Coccidioides* presence even after 15 years.

While burrows may provide favorable microhabitat for Coccidioides growth, independent of rodent presence, we found that controlling for soil conditions did not meaningfully change our estimated effect of rodents on *Coccidioides* presence via burrows. That is, only soil moisture was a meaningful predictor of *Coccidioides* presence, but mediation analyses that directly considered soil moisture as a mediator found that only 8% of the effect of burrows was mediated via altered soil moisture. Thus, our mediation analyses suggest that lowered soil moisture in burrows played only a small role in explaining why burrows might have higher probability of *C. immitis* detection, and that the within-burrow microhabitat supports *Coccidioides* via soil characteristics other than moisture, temperature, and vegetation levels – potentially humidity, soil texture and porosity, and higher concentration of key nutrients. In addition to these abiotic factors, burrows may harbor fewer microbial competitors, such as bacteria belonging to the *Bacillus* genus and fungi in the *Fennellomyces* and *Ovatospora* genera, which have been found to limit *Coccidioides* growth in laboratory experiments (Lauer, Baal et al. 2019, Kollath, Morales et al. 2023). Investigating these more specific biotic and abiotic conditions that facilitate *Coccidioides* growth in burrows will help identify areas of greatest risk for human exposure.

Associations between rodents and *Coccidioides* presence in soils may further explain observed relationships between human coccidioidomycosis incidence, precipitation, and drought. Human incidence declines markedly during drought and rebounds following heavy precipitation in years following a drought (Head, Sondermeyer-Cooksey et al. 2022). In the southwestern US, rodent host abundance also declines markedly during drought and rebounds sharply after heavy rainfall that follows the drought’s end (Prugh, Deguines et al. 2018). Post-drought rodent rebound has been linked to outbreaks of other rodent-borne infections, such as hantavirus disease (Hjelle and Glass 2000, Acuna-Soto, Stahle et al. 2002).

Our study also motivates further investigation into the role of rodent burrows on human transmission as we identified them as key sources of *Coccidioides* growth in the environment. Outbreaks of coccidioidomycosis have been observed among individuals at sites where burrows are reported (Werner and Pappagianis 1973), and among individuals excavating and re-locating burrows (Baksh, Demeter et al. 2019). This motivates protection of workers in endemic regions digging in soils that contain burrowing mammals. However, it is unknown if individuals who pass by, but do not directly disturb burrow soils, are at elevated risk of exposure to *Coccidioides* spores. A laboratory experiment buried lungs of mice infected with *C. posadasii* at the bottom of soil columns, and detected discharge of arthroconidia from the surface four weeks after inoculation, even without soil disruption (Kollath and Barker 2022). Intensive sampling around the entrance and on the top of burrows might clarify how *Coccidioides* originating within burrows might become airborne. This investigation may be particularly urgent in regions of California, where efforts are underway to phase agricultural land out of production, and there is concern that return to more natural lands – that could harbor burrowing mammal communities – may lead to more *Coccidioides* in the soil (Ellen Hanak, Alvar Escriva-Bou et al. 2019).

This study has additional limitations. The DNA assay used to detect *C. immitis* while considered the standard in the field, can only detect presence and absence of DNA, rather than quantify the amount of DNA in a sample nor the viability of the sample. *Coccidioides* has highly localized distributions in soil such that the absence of *Coccidioides* DNA in a sub-sample does not mean *Coccidioides* was not present nearby. Our statistical models measured and compared the probability of detecting *Coccidioides* DNA within a sample, and the coefficients on our statistical models reflect relative differences in average probability of detection across our sample types. In our sensitivity analyses (Supplement), we find that results are qualitatively and quantitatively similar under different classifications of a positive result. Further, detection of non-viable *Coccidiodies* DNA rather than viable *Coccidioides* spores would be expected to bias our results of the effect of rodents and their burrows towards the null.

## Conclusion

We find strong independent effects of both burrow creation and occupation on the presence of *C. immitis* in soils. Our results suggest that burrowing hosts play a critical role in shaping the distribution of *C. immitis* in soils, with much of the effect attributable to burrow creation. Increased *Coccidioides* presence in burrows likely enhances transmission cycles among rodents and may shape its transmission to humans. Our study identifies the importance of both animal hosts and their habitats – as well as their interaction – in the distribution of a human-pathogenic fungi. The methods and findings from our study could guide investigation into the synergy between reservoir host presence and soil conditions for other environmentally acquired fungal pathogens.

## Supplemental materials for

### Supplemental Methods

#### Laboratory analysis of soils

Soil samples were stored in centrifuge tubes at room temperature until analysis. We extracted DNA using the Qiagen PowerSoil kit, following established protocol (Greene, Koenig et al. 2000, Wagner, Montoya et al. 2022), and used the CocciEnv qPCR assay (Bowers, Parise et al. 2018) to determine presence of *C. immitis* in the sample. Details of the protocol are included in the Supplement. The CocciEnv assay contains 29 primer variants that target a repeating transposon sequence unique to the *Coccidioides* genome (NCBI BioProject PRJNA46299) (Bowers, Parise et al. 2018). We ran reactions in 8-well strip tubes (96 well plates) at 20μl per reaction. Each reaction contained 2μl CocciEnv primer mix, which included the 29 primers and Taqman probe (Applied Biosystems, Waltham, MA, USA) in the concentrations outlined in Bowers, et al (Bowers, Parise et al. 2018); 10μl of TaqMan Environmental Master Mix 2.0 (Applied Biosystems, Waltham, MA, USA), 6μl nuclease-free H_2_O; and 2μl of template DNA standardized to 50ng·µl^-1^. The reactions were carried out on a Stratagene Mx3000P qPCR system (Agilent Technologies, Santa Clara, CA, USA) or a Biorad-CFX96 qPCR system (BioRad Laboratories, Inc). We ran an initial cycle at 95°C for 10 minutes, followed by 40 cycles alternating between 95°C for 15 seconds and 60°C for 1 minute. We used nuclease-free water as a negative control and DNA extracted from cultured *C. posadasii* strain Silveira as a positive control. We created the standard curve from the positive control by running positive control at 7 consecutive 10-fold dilutions (highest concentration 100 ng/µL) on each plate. Amplification curves were calculated using Mxpro version 4.1 (Agilent Technologies, Santa Clara, CA, USA) with the relative fluorescence unit threshold set at 1000.

We classified an individual well as positive for *Coccidioides* if the cycle threshold (CT) was below 40, with logarithmic amplification, and the standards and negative controls behaved as expected. As qPCR amplification can produce false positives in the absence of template DNA (Ruiz-Villalba, van Pe;t-Verkuil et al. 2017), we ran four replicates of each DNA aliquot, and considered a sample was positive if it was detected in at least three of the four replicates. We conducted sensitivity analyses that varied this criteria.

*Causal mediation analysis: effect of rodents on* Coccidioides *presence, mediated by burrows* We conducted a simulation study to estimate the overall difference in the probability of *Coccidioides* detection in an area with contemporary rodent presence compared to the counterfactual scenario where rodents have never been present in that location. Following the framework developed by Wang and Arah (Wang and Arah 2015), we conducted a causal mediation analysis with g-computation, which allows us to compute the percentage of this difference that is attributable to burrow creation by rodents. We constructed directed acyclic graphs (DAG) representing the hypothesized data generating processes (Figure 3) including only rodents and burrows (Figure 3A) and adding soil conditions (Figure 3B) to allow for some of the effect of rodents and burrows to occur via soil conditions (e.g., soil moisture, temperature, vegetation). Rodents are the binary exposure of interest (*X*; 1 = present, 0 = absent), burrows the binary mediator (*M*; 1 = burrow, 0 = surface soil), and *Coccidioides* presence in soil the binary outcome (*Y*). Where used, soil conditions (moisture, temperature, vegetation) are represented among the set of covariates (***Z***), along with other key variables (sampling pasture and season).

First, we modelled the mediator conditional on the exposure and covariates, *E*(*M*|*X* = *x*, ***Z*** = ***z***); termed *m*_*x*_, where *m*_*x*1_ = *E*(*M*|*X* = 1, ***Z*** = ***z***). In this counterfactual simulation, *X* = 0 represents rodents never being present (either currently or historically) and is assumed to be approximated in the dataset as samples collected from the exclosure. The rationale for this approximation is that it allows us to satisfy the required positivity assumption for causal inference, which states that there is a nonzero probability of receiving every level of exposure for every combination of values of exposures and confounders that exist (Hernán and Robins 2006). Here, we satisfy the positivity assumption as the experimental exclosures permit every combination of the values of *M* (0/1) and *X* (0/1) to be observed. However, in the real world, *M* ≠ 1 under the scenario where *X* = 0 (i.e., burrows cannot exist where rodents have never been). Thus, instead of using our data to solve for the expected value of *M*, we deterministically set *E*(*M*|*X* = 0) = *m*_*x*0_ = 0 (i.e., rodent absence leads to burrow absence) and *E*(*M*|*X* = 1) = *m*_*x*1_ = 0.5 (i.e., rodent presence leads to equal probability of burrow or surface soil). By setting this equation deterministically, we continue to satisfy the positivity assumption required to model the expected value of the outcome (see next paragraph), while allowing the surface soil samples from the exclosure to serve as the reference group in calculation of the total effect of rodents on *Coccidioides* presence.

The outcome was modeled as dependent on the exposure, mediator, and covariates, *E*(*Y*|*X* = *x*, *M* = *m*, ***Z*** = ***z***), using the same generalized linear mixed model (GLMM) described above. As before, because soil conditions, contained within ***Z***, lie along the causal pathway, we conducted analyses that did and did not control for soil moisture, vegetation, and soil temperature. Effect estimates derived from analyses that do not control for such covariates capture the full effects of rodents and burrows. Adding soil conditions to the mediation model permits examination of the total, direct, and indirect effect of rodents on *Coccidioides* distribution, while absorbing variation that can be attributed to modified soil conditions within rodent-occupied areas (vs. rodent-excluded areas) and within burrows (vs. surface soils). Only soil moisture was a meaningful predictor of *Coccidioides* presence, so models controlling for soil conditions included only this variable.

Next, we simulated potential mediators and outcomes using our two fitted equations for *Y* and *M* (Wang and Arah 2015). Briefly, we drew 1,000 bootstrapped copies of the original sample, clustered by the highest level of clustering, which was plot (n=20). We simulated values for covariates, ***Z***, and rodent presence, *X*, that followed the same distribution as the bootstrapped data. We simulated each value of *M* by drawing from a binomial distribution centered on the deterministic specification for *m*_*x*_described above, conditional on *X*. Finally, we simulated sets of *Y*(C*occidioides* detection) using the fit GLMM, and various combinations of *x* (set to the observed distribution; set to 1; set to 0) and *m*_*x*_. For example, Y values used to determine total effect were generated using values of *x* simulated according to its observed sampling distribution and values of *m*_*x*_ solved for using the observed distribution of *x* (Table 3). We regressed the relevant set of *Y* against simulated *x* to obtain different effects (Table 3). Since our models assumed logit link functions for a binary outcome, we exponentiated results to obtain odds ratios as the effects of interest. To determine the proportion of the association between rodents and *Coccidioides* presence that is mediated via burrows, we divided the total indirect effect by the total effect prior to exponentiating.

In the causal mediation analysis, the total effect compares the probability of *Coccidioides* detection across the exposure of interest (e.g., current rodent presence vs. the counterfactual scenario of no history of rodent presence), while allowing the mediator (burrows) to take its natural value under each exposure level. The total effect can be decomposed into direct and indirect effects and is equal to the product of the pure direct effect (PDE) and the total indirect effect (TIE) (Wang and Arah 2015). The direct, unmediated effects of rodents on *Coccidioides* captures the effect of rodents that acts via pathways other than burrow creation or use. The pure direct effect (PDE) compares the probability of *Coccidioides* detection when rodents are present (*X* = 1) compared to absent (*X* = 0), under the mediator level set to the natural value it would have attained under the counterfactual, no rodent scenario (*m*_*x*0_, deterministically set to 0; Table 3).

The effect of rodents on *Coccidioides* that is explained by burrow creation (including historical) and contemporary burrow use is represented by the indirect effect. The TIE expresses the degree to which rodents cause *Coccidioides* to be present via their association with burrow presence (Figure 3A, Table 3). Mathematically, the TIE compares the probability of *Coccidioides* detection when the mediator levels are set to *m*_*x*1_(here, deterministically set to 0.5) versus *m*_*x*0_ (deterministically set to 0), while setting the exposure level to *X* = 1 (rodents present).

#### Causal mediation analysis: effect of burrows on Coccidioides presence, mediated by soil moisture

Because it has been speculated that the association between burrows and *Coccidioides* might be attributable to favorable soil conditions induced by burrows, we also examined the association between soil conditions, burrows, and *Coccidioides*. We used similar modeling framework as described above, with each soil condition (i.e., soil moisture, vegetation, temperature) considered as the mediator in turn and burrows or surface soils the exposure (Figure S7). The DAG representing the hypothesized data generating processes is included in Figure S7. Burrows are the binary exposure of interest (*X*; 1 = burrow, 0 = surface soil), soil conditions the mediator (continuous), and *Coccidioides* presence in soil the binary outcome (*Y*). Rodents are represented among the set of covariates (***Z***), along with other key variables (sampling season).

All steps for this mediation analysis were similar to the above, except for this analysis, we could model the mediator conditional on the exposure and covariates (*m*_*x*_) using observed data. We did so using a generalized linear mixed model with fixed effects on burrows (*X*) and the listed covariates (***Z***; rodents and season), and random effects on sampling cluster nested within plot. Because soil moisture is a continuous variable, we assumed a gaussian distribution, and sampled values for the mediator from a normal distribution with mean centered at the model predictions and standard deviation as ascertained from the model. The GLMM that solved for the expected value of the outcome conditional on the exposure and mediators was updated to include the mediator of interest. Because soil moisture was the only measured characteristic found to be associated with *Coccidioides* presence, we report results only for soil moisture as the mediator.

### Supplemental Results

#### Clustering of positive samples

We observed significant clustering of samples at the precinct and the transect level. Among burrow samples collected, samples collected from the same precinct were more likely to be similar to each other than under a random distribution that does not account for clustering (Figure S5). Among samples collected from outside the exclosures, we observed that 14.5% of the five-sample clusters belonging to the same precinct contained four or more positive samples. Using the cumulative density function of a binomial distribution with five trials and probability of success equal to 0.285 (the proportion of samples from burrows with rodents that were positive), we would expect that only 2.5% of the five-sample clusters to contain four or more positive samples. Among samples collected from inside the exclosures, we observed that 11.4% of the five-sample clusters belonging to the same precinct contained four or more positive samples. Using the cumulative density function of a binomial distribution with five trials and probability of success equal to 0.197 (the proportion of samples from burrows without rodents that were positive), we would expect that only 0.6% of the five-sample clusters to contain four or more positive samples.

We identified *Coccidioides* in 15 of the 790 surface soil samples, originating from eight transects. Clustering of positive surface soil samples from outside the exclosure was observed: three transects each yielded two positive samples in the same season and another four transects yielded two positive samples across multiple seasons. Among samples collected from outside the exclosures, we observed that 3.8% of the five-sample clusters belonging to the same transect contained two or more positive samples. Using the cumulative density function of a binomial distribution with five trials and probability of success equal to 0.035, we would expect that only 1.1% of the five-sample clusters to contain two or more positive samples. There was no statistically significant association between *Coccidioides* detection and distance away from the precinct, nor was there an association between positivity among the transects and positivity in the precinct.

#### Sensitivity analyses

When we varied the definition of a positive sample for *Coccidioides* to be between one and four replicates with a CT value equal to or below 40 (from our primary definition of three of four replicates), the total number of samples positive for *Coccidioides* varied from 13.3% (266 samples with four of four replicates with CT value under 40) to 21.7% (432 samples with at least one of four replicates with CT value under 40; Table S3). Overall, conclusions regarding the relationships between rodents, burrows, and *Coccidioides* detection did not change when we varied the definition of a positive sample (Tables S4-S7). Using a positive sample definition of one of four replicates yielded lower coefficients on burrows, and a lower proportion of the total effect that was mediated via burrows (i.e., a larger percentage of the total effect was direct). Several exclosures exhibited potential evidence of recent breaching. While we did not collect soils from any burrows that appeared active within the exclosure, we repeated our mediation analysis dropping two plots where concern of breaching had been noted across more than one sampling season. In one plot, the fence wall had partially collapsed, reducing the height of the fence to about 1 foot, which could have allowed rodents to climb over; in the other, the fence appeared intact, but several burrows exhibited signs of recent activity (e.g., feet or tail markings in the dirt). We hypothesized that rodent breachings would make the exclosures appear more like non-exclosures, thereby artificially reducing the effect of rodents on *Coccidioides* presence towards the null. When we excluded these experimental plots from the analysis, the estimated coefficients from the GLMM and the mediation analysis were similar, although the direct effect of rodents decreased slightly and the proportion of the total effect of rodents on *Coccidioides* presence that was mediated by burrows slightly increased (Tables S4-S7). Accordingly, the bias was not in the direction suspected, which reduces concerns about the influence of imperfect exclosures.

---

**Figure S1.**
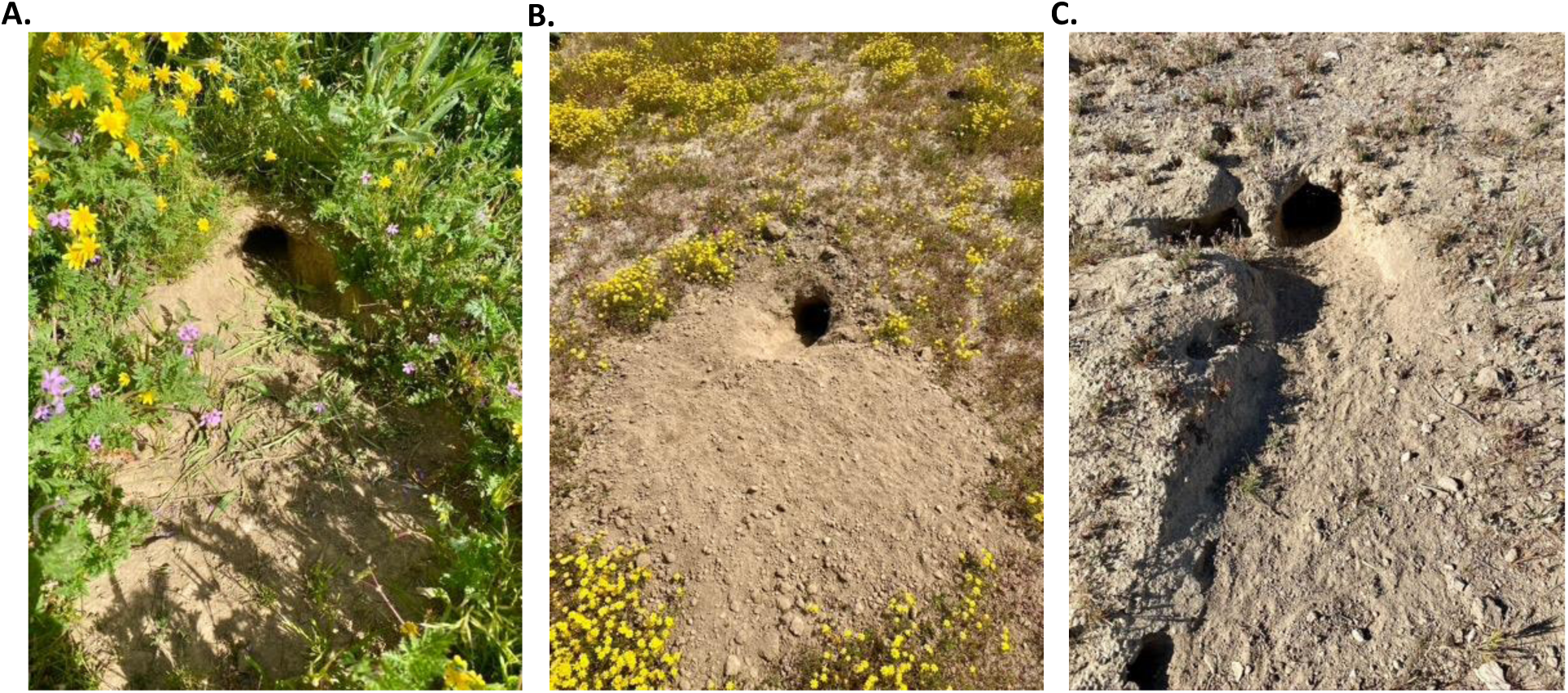
**Images of burrow entrances showing signs of giant kangaroo rat activity**, including clipped vegetation (A) and tail and feet markings (B,C). Photographs taken by Simon K. Camponuri.

**Figure S2.**
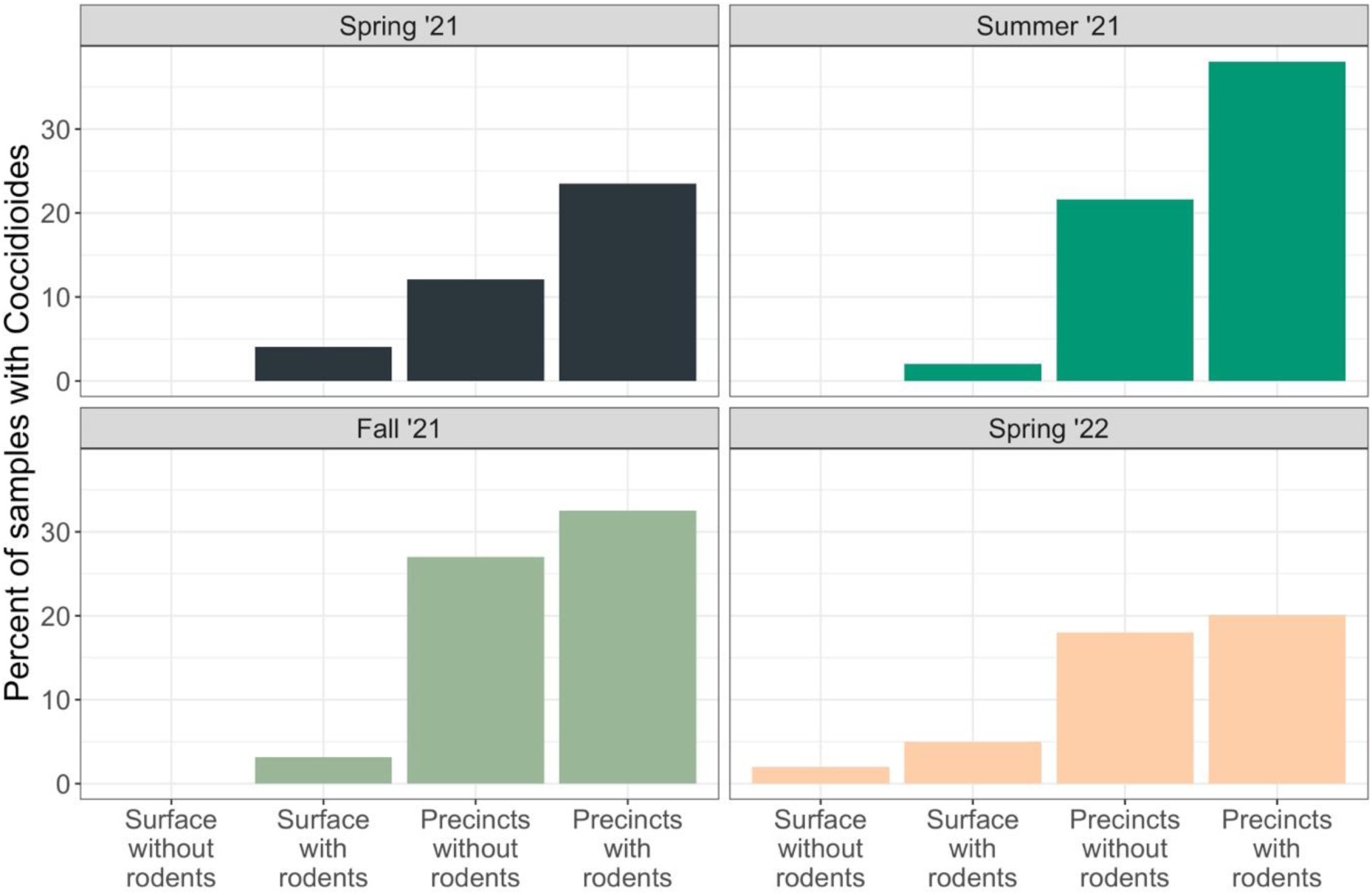
Probability (%) of *C. immitis* detection by season and sample type.

**Figure S3.**
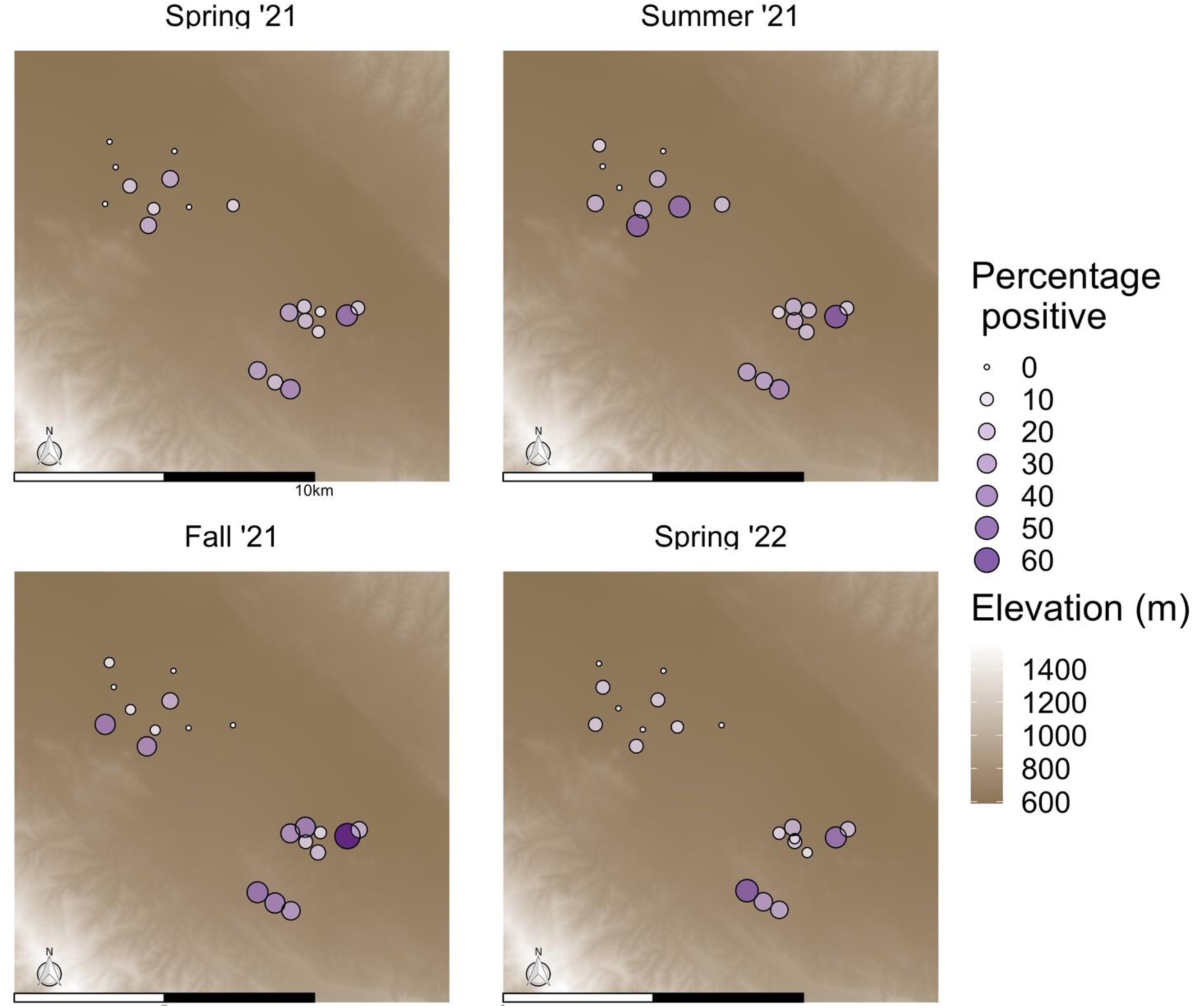
Percentage (%) of samples per plot that tested positive for Coccidioides in soils, stratified by season of sampling. Each symbol represents all samples drawn from one experimental plot in the Carrizo Plain Ecosystem Project, within each sampling seasons (n = ∼25 samples per season, totaling approximately 100 per plot across four seasons).

**Figure S4.**
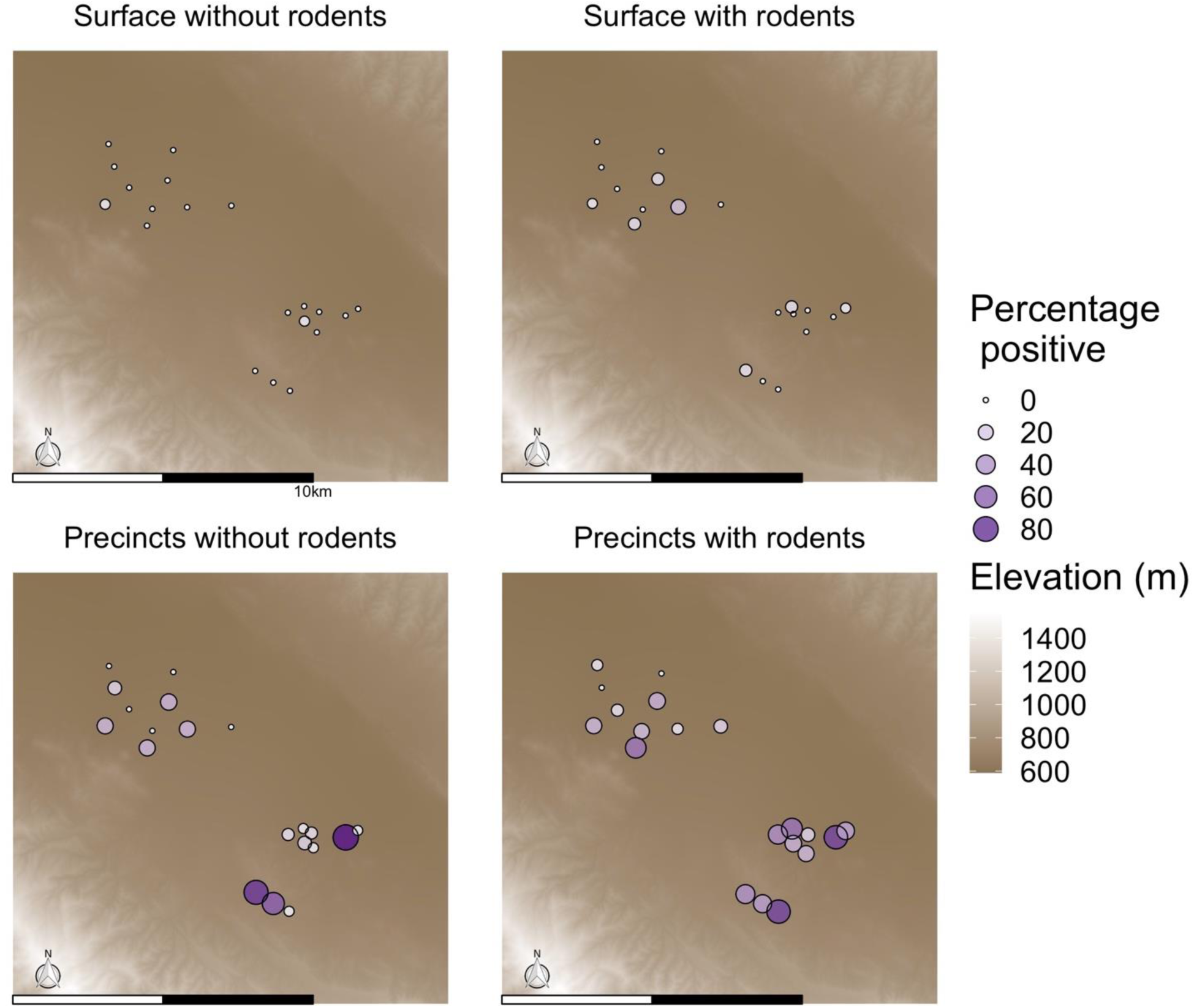
Percentage (%) of samples per plot that tested positive for Coccidioides in soils, stratified by type of sample. Each symbol represents samples of a single type drawn from one experimental plot in the Carrizo Plain Ecosystem Project, across each sampling seasons.

**Figure S5.**
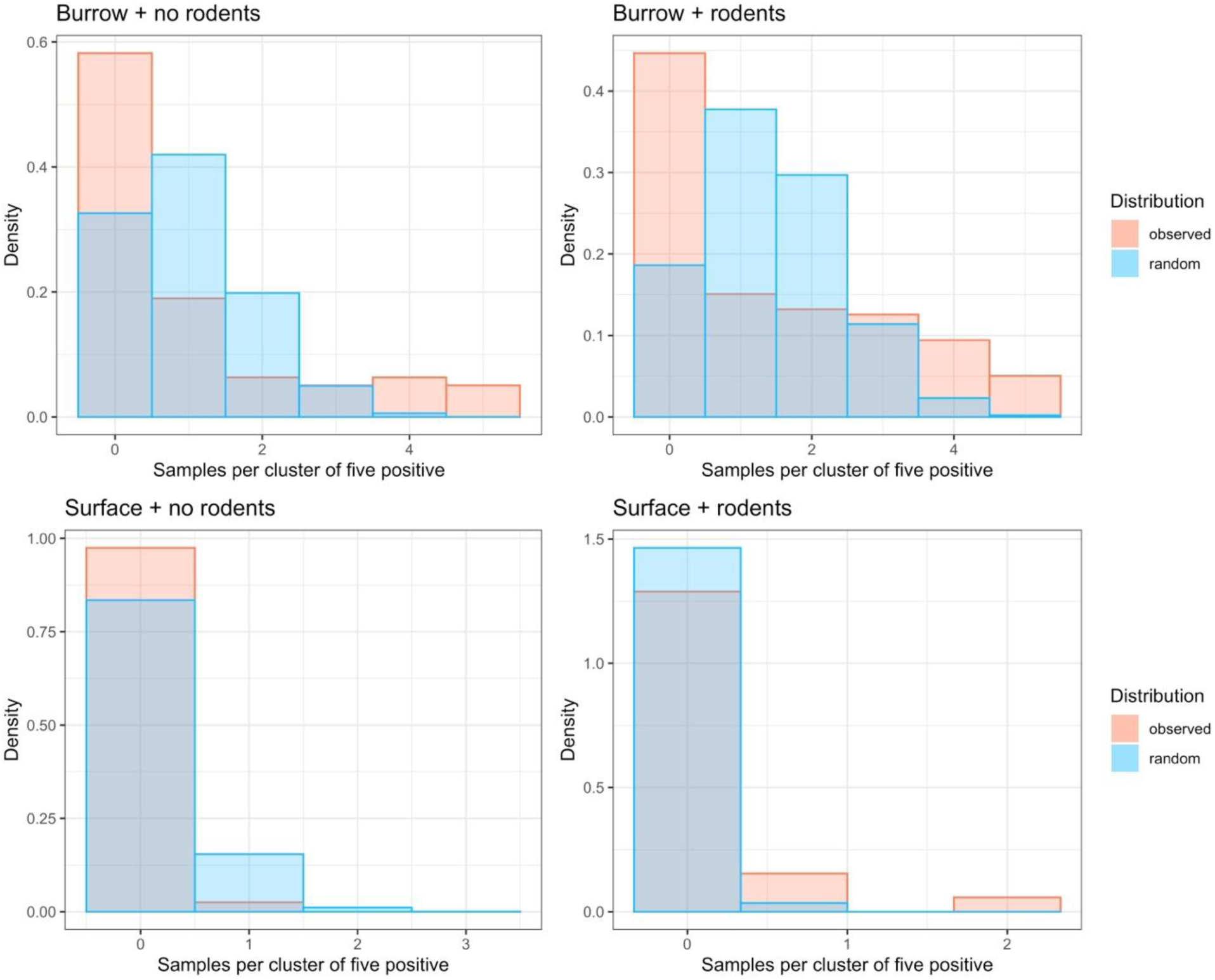
Evidence of clustering of positive samples within precincts and transects. Each cluster contains five samples collected from the same precinct (burrow) or transect (surface). Orange colors represent the density of the number of positive samples from each precinct observed. Blue colors represent the density expected under a random distribution, with no significant clustering.

**Figure S6.**
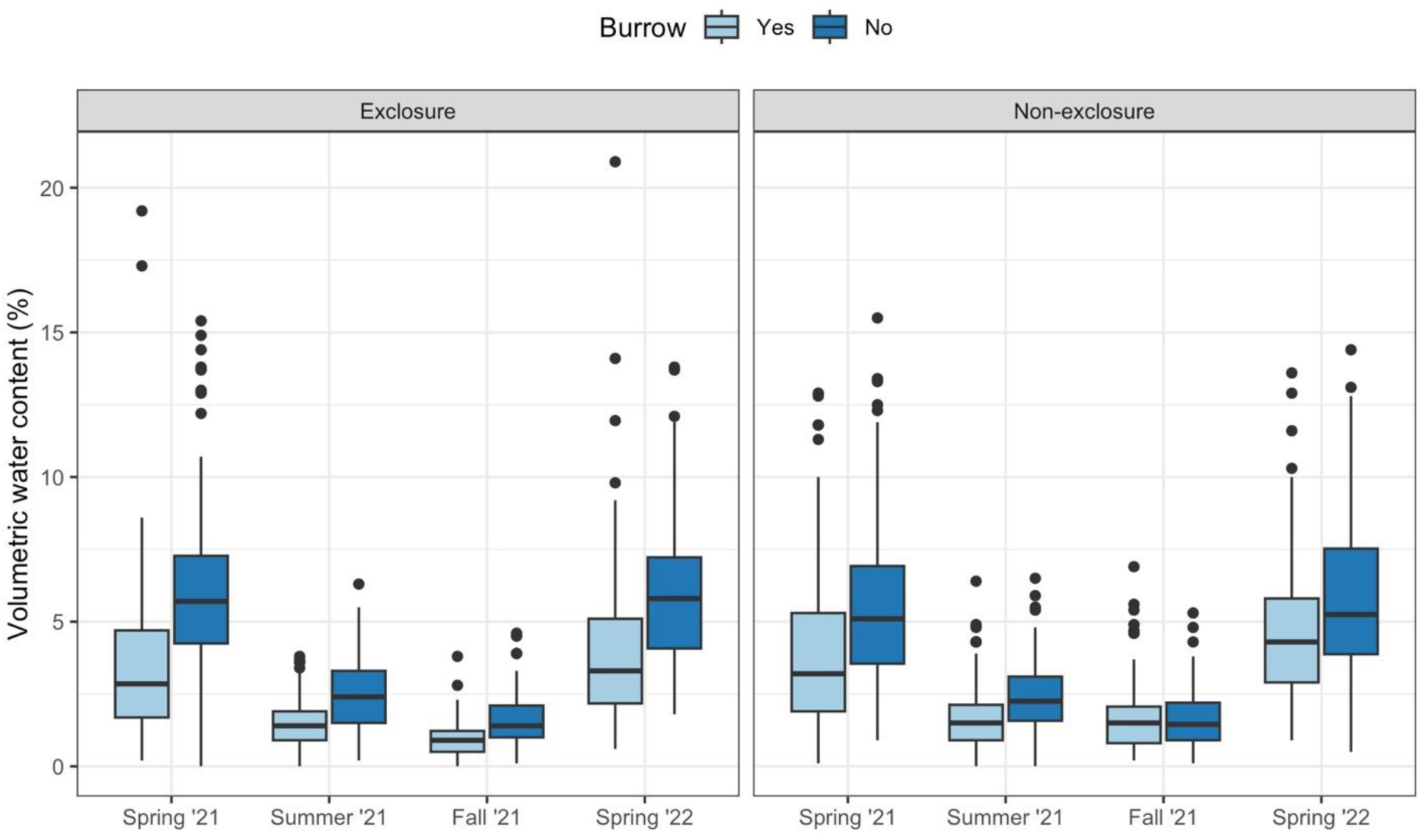
Comparison of soil moisture, measured as percent volumetric water content, across season and sample type.

**Figure S7.**
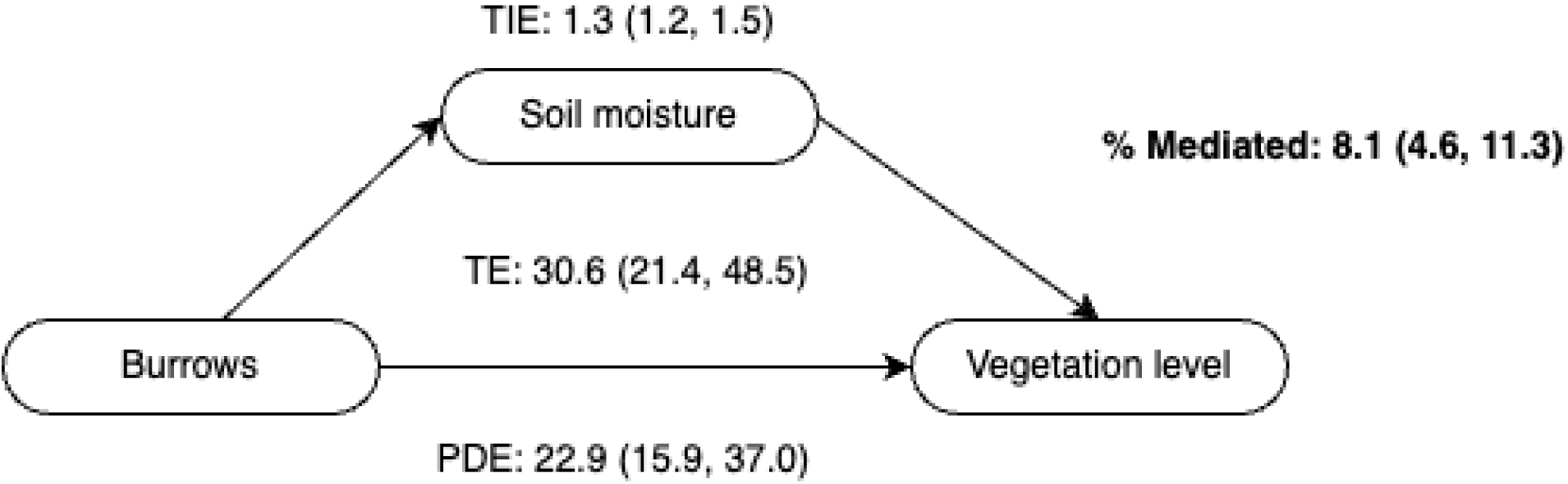
**Directed acyclic graphs (DAGs) representing the hypothesized relationships between rodents, burrows, and soil conditions**, including moisture (A), temperature (B), and vegetation level (C; ordinal). TE = total effect; TDE = total direct effect; PDE = pure direct effect; TIE = total indirect effect; PIE = pure indirect effect.

**Figure S8.**
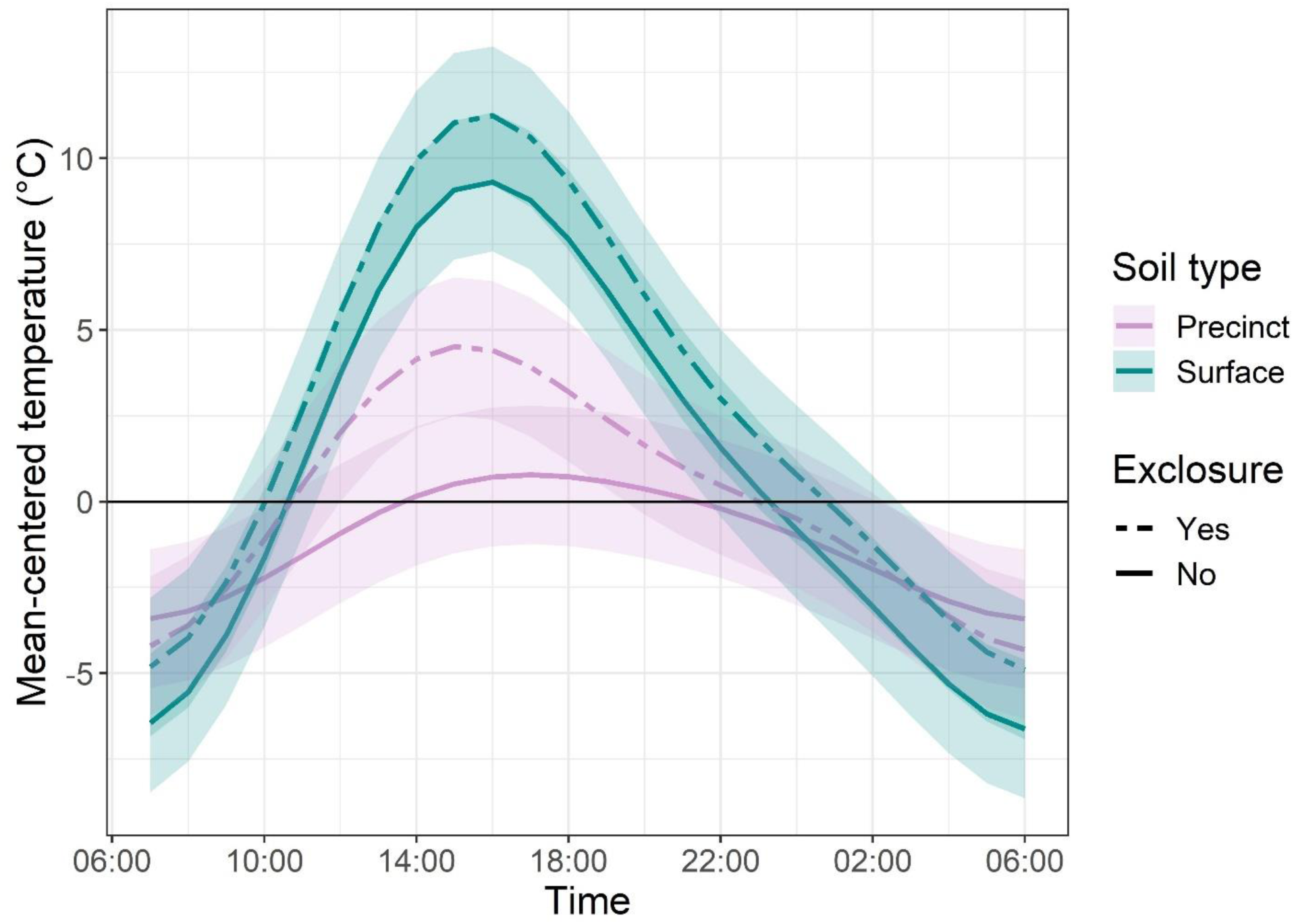
Comparison of continuous temperature measurement across burrows, surface soils, rodent exclosures, and rodent non-exclosures.

**Table S1.**
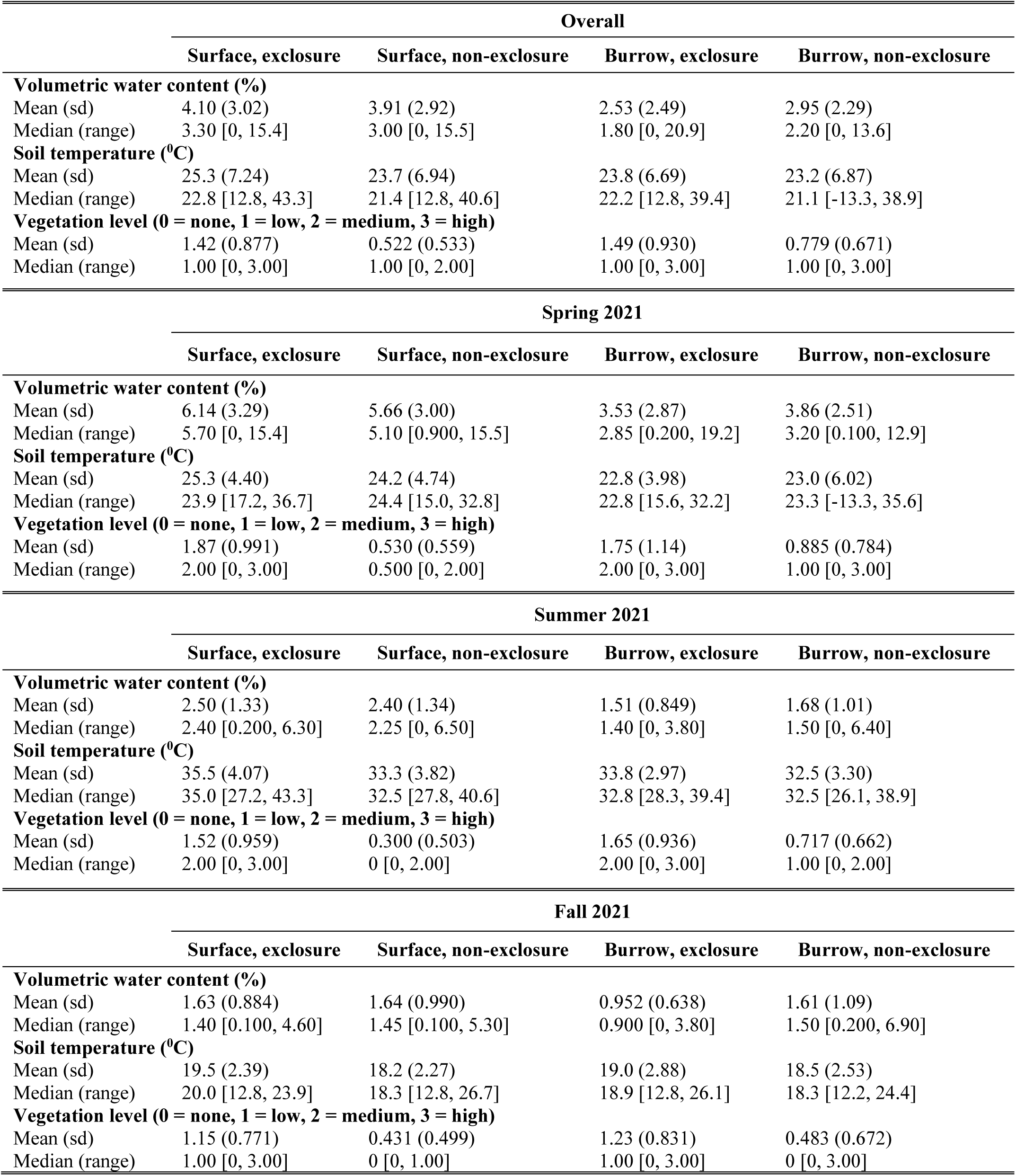
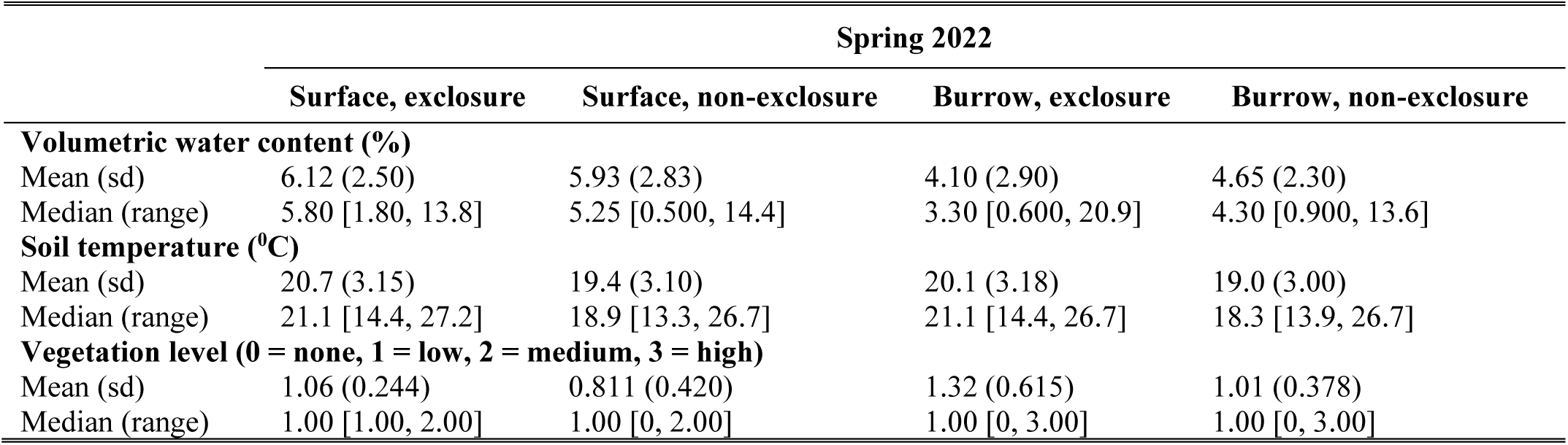
Comparison of soil moisture (expressed as volumetric water content), soil temperature, and vegetation level by season and sample type. Exclosures means rodents are not present, non-exclosure means rodents may be present.

| Overall |  |  |  |  |
| --- | --- | --- | --- | --- |
|  | Surface, exclosure | Surface, non-exclosure | Burrow, exclosure | Burrow, non-exclosure |
| **Volumetric water content (%)** |  |  |  |  |
| Mean (sd) | 4.10 (3.02) | 3.91 (2.92) | 2.53 (2.49) | 2.95 (2.29) |
| Median (range) | 3.30 [0, 15.4] | 3.00 [0, 15.5] | 1.80 [0, 20.9] | 2.20 [0, 13.6] |
| **Soil temperature (°C)** |  |  |  |  |
| Mean (sd) | 25.3 (7.24) | 23.7 (6.94) | 23.8 (6.69) | 23.2 (6.87) |
| Median (range) | 22.8 [12.8, 43.3] | 21.4 [12.8, 40.6] | 22.2 [12.8, 39.4] | 21.1 [-13.3, 38.9] |
| **Vegetation level (0 = none, 1 = low, 2 = medium, 3 = high)** |  |  |  |  |
| Mean (sd) | 1.42 (0.877) | 0.522 (0.533) | 1.49 (0.930) | 0.779 (0.671) |
| Median (range) | 1.00 [0, 3.00] | 1.00 [0, 2.00] | 1.00 [0, 3.00] | 1.00 [0, 3.00] |
| Spring 2021 |  |  |  |  |
|  | Surface, exclosure | Surface, non-exclosure | Burrow, exclosure | Burrow, non-exclosure |
| **Volumetric water content (%)** |  |  |  |  |
| Mean (sd) | 6.14 (3.29) | 5.66 (3.00) | 3.53 (2.87) | 3.86 (2.51) |
| Median (range) | 5.70 [0, 15.4] | 5.10 [0.900, 15.5] | 2.85 [0.200, 19.2] | 3.20 [0.100, 12.9] |
| **Soil temperature (°C)** |  |  |  |  |
| Mean (sd) | 25.3 (4.40) | 24.2 (4.74) | 22.8 (3.98) | 23.0 (6.02) |
| Median (range) | 23.9 [17.2, 36.7] | 24.4 [15.0, 32.8] | 22.8 [15.6, 32.2] | 23.3 [-13.3, 35.6] |
| **Vegetation level (0 = none, 1 = low, 2 = medium, 3 = high)** |  |  |  |  |
| Mean (sd) | 1.87 (0.991) | 0.530 (0.559) | 1.75 (1.14) | 0.885 (0.784) |
| Median (range) | 2.00 [0, 3.00] | 0.500 [0, 2.00] | 2.00 [0, 3.00] | 1.00 [0, 3.00] |
| Summer 2021 |  |  |  |  |
|  | Surface, exclosure | Surface, non-exclosure | Burrow, exclosure | Burrow, non-exclosure |
| **Volumetric water content (%)** |  |  |  |  |
| Mean (sd) | 2.50 (1.33) | 2.40 (1.34) | 1.51 (0.849) | 1.68 (1.01) |
| Median (range) | 2.40 [0.200, 6.30] | 2.25 [0, 6.50] | 1.40 [0, 3.80] | 1.50 [0, 6.40] |
| **Soil temperature (°C)** |  |  |  |  |
| Mean (sd) | 35.5 (4.07) | 33.3 (3.82) | 33.8 (2.97) | 32.5 (3.30) |
| Median (range) | 35.0 [27.2, 43.3] | 32.5 [27.8, 40.6] | 32.8 [28.3, 39.4] | 32.5 [26.1, 38.9] |
| **Vegetation level (0 = none, 1 = low, 2 = medium, 3 = high)** |  |  |  |  |
| Mean (sd) | 1.52 (0.959) | 0.300 (0.503) | 1.65 (0.936) | 0.717 (0.662) |
| Median (range) | 2.00 [0, 3.00] | 0 [0, 2.00] | 2.00 [0, 3.00] | 1.00 [0, 2.00] |
| Fall 2021 |  |  |  |  |
|  | Surface, exclosure | Surface, non-exclosure | Burrow, exclosure | Burrow, non-exclosure |
| **Volumetric water content (%)** |  |  |  |  |
| Mean (sd) | 1.63 (0.884) | 1.64 (0.990) | 0.952 (0.638) | 1.61 (1.09) |
| Median (range) | 1.40 [0.100, 4.60] | 1.45 [0.100, 5.30] | 0.900 [0, 3.80] | 1.50 [0.200, 6.90] |
| **Soil temperature (°C)** |  |  |  |  |
| Mean (sd) | 19.5 (2.39) | 18.2 (2.27) | 19.0 (2.88) | 18.5 (2.53) |
| Median (range) | 20.0 [12.8, 23.9] | 18.3 [12.8, 26.7] | 18.9 [12.8, 26.1] | 18.3 [12.2, 24.4] |
| **Vegetation level (0 = none, 1 = low, 2 = medium, 3 = high)** |  |  |  |  |
| Mean (sd) | 1.15 (0.771) | 0.431 (0.499) | 1.23 (0.831) | 0.483 (0.672) |
| Median (range) | 1.00 [0, 3.00] | 0 [0, 1.00] | 1.00 [0, 3.00] | 0 [0, 3.00] |

| Spring 2022 |  |  |  |  |
| --- | --- | --- | --- | --- |
|  | Surface, exclosure | Surface, non-exclosure | Burrow, exclosure | Burrow, non-exclosure |
| **Volumetric water content (%)** |  |  |  |  |
| Mean (sd) | 6.12 (2.50) | 5.93 (2.83) | 4.10 (2.90) | 4.65 (2.30) |
| Median (range) | 5.80 [1.80, 13.8] | 5.25 [0.500, 14.4] | 3.30 [0.600, 20.9] | 4.30 [0.900, 13.6] |
| **Soil temperature (<sup>0</sup>C)** |  |  |  |  |
| Mean (sd) | 20.7 (3.15) | 19.4 (3.10) | 20.1 (3.18) | 19.0 (3.00) |
| Median (range) | 21.1 [14.4, 27.2] | 18.9 [13.3, 26.7] | 21.1 [14.4, 26.7] | 18.3 [13.9, 26.7] |
| **Vegetation level (0 = none, 1 = low, 2 = medium, 3 = high)** |  |  |  |  |
| Mean (sd) | 1.06 (0.244) | 0.811 (0.420) | 1.32 (0.615) | 1.01 (0.378) |
| Median (range) | 1.00 [1.00, 2.00] | 1.00 [0, 2.00] | 1.00 [0, 3.00] | 1.00 [0, 3.00] |

**Table S2.**
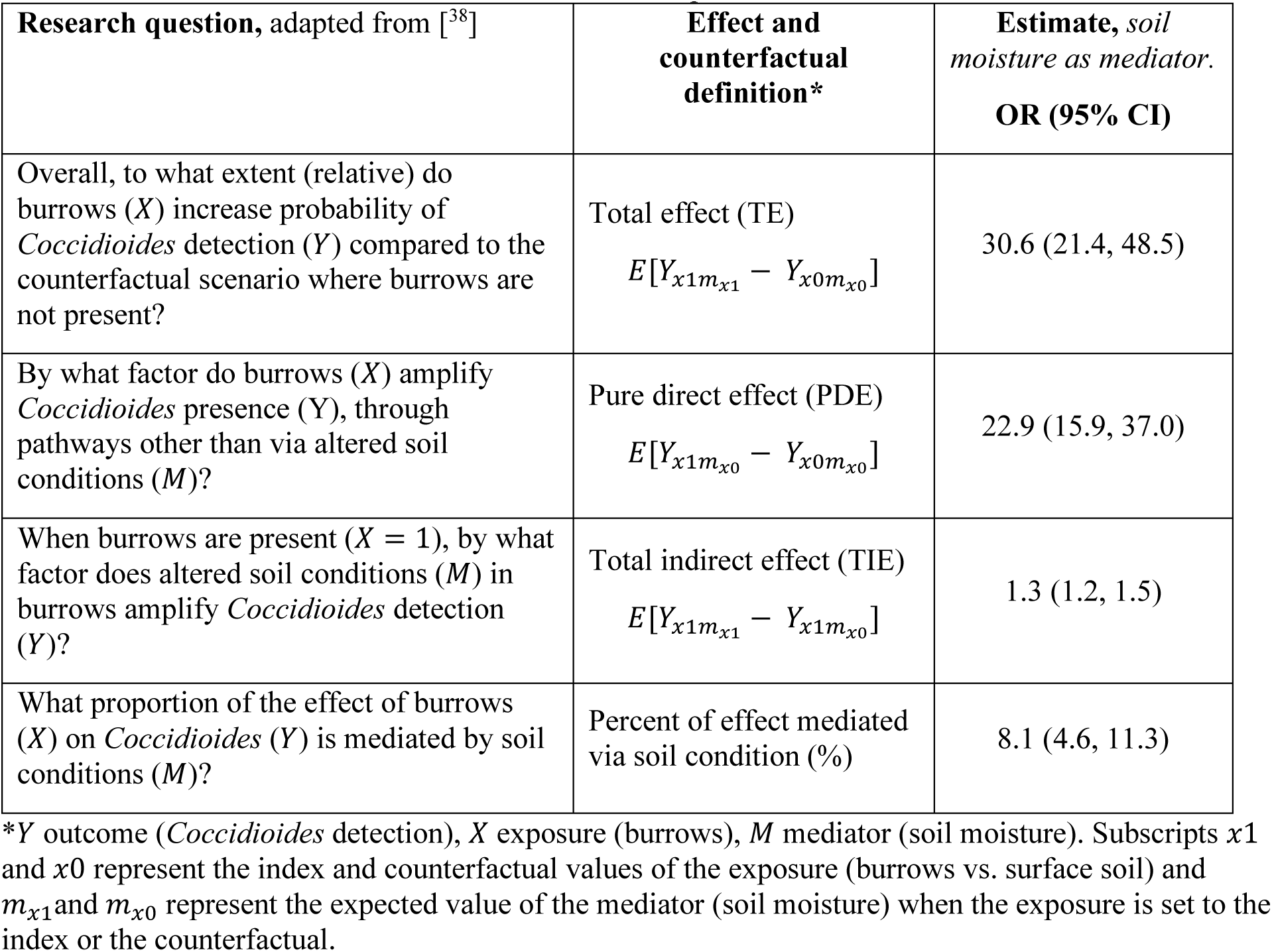
Odds ratios representing various effects from causal mediation analysis examining relationship between burrows, soil moisture, and C. immitis in soils. Here, burrows are the exposure of interest, soil moisture the mediator, and *Coccidioides* present the outcome.

**Table S3.** Prevalence of Coccidioides detection by sample type, using differing classification cutoffs for positivity. The main analysis uses a cut-off of 3 positives per 4 replicates.

| Replicate<br>wells with CT<br>value $\leq 40$ | Burrows | | Surface soils | | Total |
| --- | --- | --- | --- | --- | --- |
|  | Rodents excluded | Rodents present | Rodents excluded | Rodents present |  |
| **1 of 4** | 96/396 (24.2%) | 300/799 (37.5%) | 10/399 (2.5%) | 26/394 (6.6%) | 432/1988 (21.7%) |
| **2 of 4** | 86/396 (21.7%) | 262/799 (32.8%) | 3/399 (0.75%) | 18/394 (4.5%) | 369/1988 (18.5%) |
| **3 of 4 (main)** | 78/396 (19.7%) | 228/799 (28.5%) | 2/399 (0.5%) | 14/394 (3.6%) | 322/1988 (16.2%) |
| **4 of 4** | 70/396 (17.7%) | 182/799 (22.8%) | 2/399 (0.5%) | 12/394 (3.1%) | 266/1988 (13.3%) |

**Table S4.**
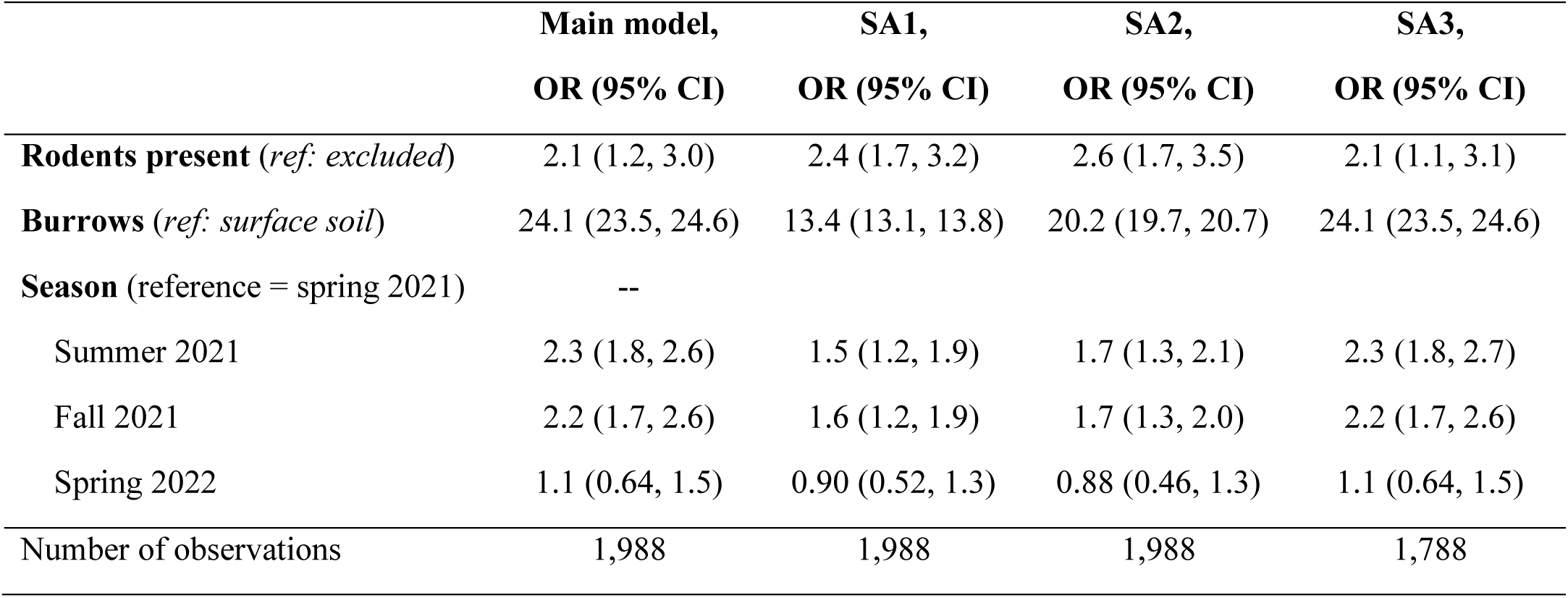
Sensitivity analyses displaying results from the generalized linear mixed model, not controlling for soil conditions. SA1: Sensitivity analyses where a positive sample required only one of the 4 replicates to have a CT value less than 40. SA2: Sensitivity analysis where a positive sample required only two of the 4 replications to have a CT value less than 40. SA3: Excluding two plots (one from each pasture) where rodent exclosure was noted to have potential rodent activity during two or more sampling trips. Main analysis includes all 20 plots and considers a positive detection to have three of 4 replicates with a CT value less than 40. These results are also in Table 2 of the main manuscript.

|  | **Main model,** | **SA1,** | **SA2,** | **SA3,** |
| --- | --- | --- | --- | --- |
|  | **OR (95% CI)** | **OR (95% CI)** | **OR (95% CI)** | **OR (95% CI)** |
| **Rodents present** (*ref: excluded*) | 2.1 (1.2, 3.0) | 2.4 (1.7, 3.2) | 2.6 (1.7, 3.5) | 2.1 (1.1, 3.1) |
| **Burrows** (*ref: surface soil*) | 24.1 (23.5, 24.6) | 13.4 (13.1, 13.8) | 20.2 (19.7, 20.7) | 24.1 (23.5, 24.6) |
| **Season** (reference = spring 2021) | -- |  |  |  |
| Summer 2021 | 2.3 (1.8, 2.6) | 1.5 (1.2, 1.9) | 1.7 (1.3, 2.1) | 2.3 (1.8, 2.7) |
| Fall 2021 | 2.2 (1.7, 2.6) | 1.6 (1.2, 1.9) | 1.7 (1.3, 2.0) | 2.2 (1.7, 2.6) |
| Spring 2022 | 1.1 (0.64, 1.5) | 0.90 (0.52, 1.3) | 0.88 (0.46, 1.3) | 1.1 (0.64, 1.5) |
| Number of observations | 1,988 | 1,988 | 1,988 | 1,788 |

**Table S5.** Sensitivity analyses displaying results from the generalized linear mixed model, controlling for soil conditions. SA1: Sensitivity analyses where a positive sample required only one of the 4 replicates to have a CT value less than 40. SA2: Sensitivity analysis where a positive sample required only two of the 4 replications to have a CT value less than 40. SA3: Excluding two plots (one from each pasture) where rodent exclosure was noted to have potential rodent activity during two or more sampling trips. Main model includes all 20 plots and considers a positive detection to have 3 of 4 replicates with a CT value less than 40. These results are also in Table 2 of the main manuscript.

|  | **Main model,** | **SA1,** | **SA2,** | **SA3,** |
| --- | --- | --- | --- | --- |
|  | **OR (95% CI)** | **OR (95% CI)** | **OR (95% CI)** | **OR (95% CI)** |
| **Rodents present** (*ref: excluded*) | 2.5 (1.6, 3.4) | 2.5 (1.7, 3.3) | 2.7 (1.8, 3.5) | 2.2 (1.3, 3.2) |
| **Burrows** (*ref: surface soil*) | 20.2 (19.7, 20.8) | 12.2 (11.8, 12.6) | 18.8 (18.3, 19.3) | 20.5 (19.9, 21.1) |
| **Soil moisture** (water content %) | 0.87 (0.76, 0.98) | 0.91 (0.82, 0.99) | 0.93 (0.84, 1.0) | 0.83 (0.72, 0.95) |
| **Season** (reference = spring 2021) |  |  |  |  |
| Summer 2021 | 1.7 (1.3, 2.2) | 1.3 (0.86, 1.7) | 1.5 (1.0, 1.9) | 1.7 (1.2, 2.1) |
| Fall 2021 | 1.6 (1.1, 2.0) | 1.2 (0.85, 1.7) | 1.4 (0.98, 1.9) | 1.5 (1.0, 2.0) |
| Spring 2022 | 1.2 (0.75, 1.6) | 0.97 (0.59, 1.4) | 0.93 (0.52, 1.3) | 1.3 (0.79, 1.7) |
| Number of observations | 1,997 | 1,988 | 1,988 | 1,788 |

**Table S6.**
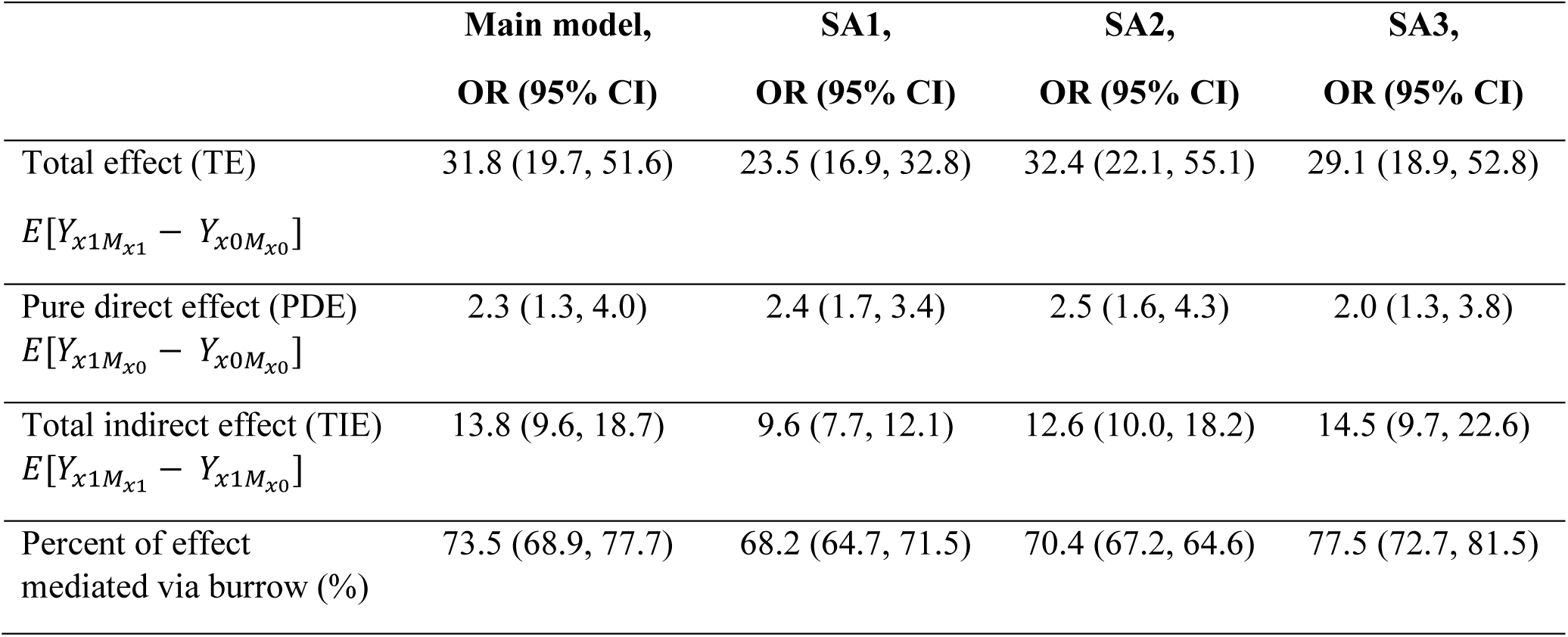
Sensitivity analyses displaying results from mediation analysis, not controlling for soil conditions. SA1: Sensitivity analyses where a positive sample required only one of the 4 replicates to have a CT value less than 40. SA2: Sensitivity analysis where a positive sample required only two of the 4 replications to have a CT value less than 40. SA3: Excluding two plots (one from each pasture) where rodent exclosure was noted to have potential rodent activity during two or more sampling trips. Main analysis includes all 20 plots and considers a positive detection to have 3 of 4 replicates with a CT value less than 40. These results are also in Table 3 of the main manuscript and in Figure 4A.

**Table S7.** Sensitivity analyses displaying results from mediation analysis, controlling for soil conditions. SA1: Sensitivity analyses where a positive sample required only one of the 4 replicates to have a CT value less than 40. SA2: Sensitivity analysis where a positive sample required only two of the 4 replications to have a CT value less than 40. SA3: Excluding two plots (one from each pasture) where rodent exclosure was noted to have potential rodent activity during two or more sampling trips. Main analysis includes all 20 plots and considers a positive detection to have 3 of 4 replicates with a CT value less than 40. These results are also in Table 3 of the main manuscript and in Figure 4B.

|  | **Main model,** | **SA1,** | **SA2,** | **SA3,** |
| --- | --- | --- | --- | --- |
|  | **OR (95% CI)** | **OR (95% CI)** | **OR (95% CI)** | **OR (95% CI)** |
| Total effect (TE)<br>$E[Y_{x1M_{x1}} - Y_{x0M_{x0}}]$ | 28.9 (18.7, 55.6) | 21.7 (16.8, 30.5) | 29.7 (19.5, 56.3) | 26.3 (16.8, 43.8) |
| Pure direct effect (PDE)<br>$E[Y_{x1M_{x0}} - Y_{x0M_{x0}}]$ | 2.5 (1.5, 4.9) | 2.4 (1.8, 3.5) | 2.6 (1.8, 4.5) | 2.3 (1.3, 4.3) |
| Total indirect effect (TIE)<br>$E[Y_{x1M_{x1}} - Y_{x1M_{x0}}]$ | 11.4 (8.4, 16.6) | 8.8 (7.2, 10.8) | 11.1 (8.7, 14.9) | 11.6 (8.2, 16.8) |
| Percent of effect mediated via burrow (%) | 70.6 (65.6, 75.4) | 66.9 (63.8, 70.7) | 69.0 (64.4, 72.9) | 74.1 (69.8, 78.5) |

